# Advancing multi-day ex vivo kidney perfusion using spatially resolved metabolomics

**DOI:** 10.1101/2023.05.10.540143

**Authors:** Marlon J.A. de Haan, Franca M.R. Witjas, Annemarie M.A. de Graaf, Marleen E. Jacobs, Elena Sánchez-López, Sarantos Kostidis, Martin Giera, Mehdi Maanaoui, Thomas Hubert, Julie Kerr-Conte, François Pattou, Dorottya K. de Vries, Jesper Kers, Ian P.J. Alwayn, Cees van Kooten, Bram P.A.M. Heijs, Gangqi Wang, Marten A. Engelse, Ton J. Rabelink

## Abstract

The ability to preserve metabolically active kidneys ex vivo for multiple days may permit reconditioning, repair and regeneration of deceased donor kidneys. However, the kidneys high metabolic demand limits its functional preservation. Current approaches focus on normothermic machine perfusion (NMP) at 37°C or hypothermic machine perfusion (HMP) at 4-8°C. At normothermia, kidneys are metabolically active but *ex vivo* preservation is limited to hours. During hypothermia kidneys can be preserved up to 24 hours but are metabolically inactive and suffer cold-induced injury. Therefore, we revisited sub normothermic perfusion (at 25°C) as an alternative approach to preserve human kidneys in a metabolically active state for extended periods of time.

In a custom-made platform that includes a cell-free perfusate enriched with TCA cycle fuels, urine recirculation, and continuous hemofiltration we perfused discarded human kidneys up to 8 days. Using spatially resolved single cell resolution isotope tracing we demonstrate active metabolism in all the different renal cell types over this period. However, beyond 4 days cell composition of nephron segments assessed with spatial lipidomics changed substantially and injury markers such as NGAL and LDH increased in the perfusate. Up to 4 days, perfused human discarded donor kidneys maintained metabolic fluxes, functional parameters and allow for reperfusion using a porcine auto transplantation model. These data underpin that extended multi-day metabolic preservation of human kidneys is achievable using a sub normothermic perfusion platform.

## INTRODUCTION

During the past decade machine perfusion has (re)gained broad interest for kidney preservation (1). It has been demonstrated that machine perfusion improves transplantation outcomes as compared to static cold storage (SCS) (2, 3) and allows the transplantation of organs that otherwise would have been discarded (4, 5). This has resulted in a paradigm shift from SCS-based preservation towards perfusion-based preservation. With technological advances and development of more complex perfusion platforms, interest in pre-transplant preservation of kidneys in a metabolically active state has sparked, with the prospect of resuscitation, repair, rejuvenation, and perhaps even regeneration (6, 7). Whereas short-term machine perfusion (≤6h) allows for resuscitation and viability assessment, extended preservation is likely needed for reconditioning, repair and regeneration of the graft (8–10).

*Ex viv*o preservation of human livers has been prolonged to 7-days (11), with promising 1-year post transplant outcomes for a patient that received a liver graft after 3-days of ex vivo preservation (7, 9, 12). In contrast to the advances seen in multi-day preservation of human livers (6, 8, 11–13), preservation of donor kidneys in a metabolically active state seems far more challenging. Reports of successful kidney perfusion up to or beyond 24h are rare (9, 14, 15).

Whereas most groups focus on normothermic perfusion (NMP) for prolonged (≥6h) kidney preservation (14, 16, 17), the use of oxygen carriers, although effective, can be hazardous for the kidney graft. Erythrocyte damage and haemolysis are common sequelae of extracorporeal perfusion platforms, resulting in RBC depletion and therewith free haemoglobin accumulation (18). Whereas the liver can convert free haemoglobin into bilirubin and excrete it into bile during multi-day perfusion (11, 19), free haemoglobin will accumulate during kidney perfusion. This poses a serious problem given that the kidney is extremely susceptible to supraphysiological levels of cell free haemoglobin, with acute kidney injury (AKI) as a common consequence (20–25). Cell free haemoglobin was identified as functional second hit that exacerbates ischemia-reperfusion-injury (IRI) to overt AKI in a mouse model (23). Therefore, we postulate that omission of oxygen carriers, and thus an acellular perfusion approach will better support multi-day kidney preservation.

Two decades ago, *Brasile et al.* proposed subnormothermic perfusion as approach for prolonged *ex vivo* preservation of donor kidneys (15, 26, 27). Lowering preservation temperature reduces the metabolic rate, and therewith oxygen requirements. It has been demonstrated that at sub normothermia (20-32°C), the oxygen demand of the graft is low enough to be satisfied through an acellular perfusate, thereby negating the need for oxygen carriers (28–31). Moreover, it was recently demonstrated that in principle sufficient metabolic activity is maintained at sub normothermia (28°C) to support molecular and cellular repair processes (32).

To this end we developed a sub normothermic (25°C) kidney culture platform that supports multi-day *ex vivo* preservation of discarded human kidneys. The kidney is an inherently complex organ with over 20 different cell types that are responsible for continuous reabsorption of solutes and metabolites against concentration gradients in a process that demands a large amount of energy (33, 34). These metabolic needs are predominantly driven by proximal tubular cells (PTs). They rely on oxidative mitochondrial metabolism to meet their energy needs (35, 36). Fatty acids, citrate and glutamine are the main nutrient sources for PTs, rather than glucose (34, 37, 38). Therefore, instead of standard electrolyte solutions containing few ingredients, alternative nutrient sources like glutamine and other TCA cycle fuels were included in the platform to support these metabolic preferences.

We sought to extend perfusion time beyond 24h and performed cell-type-specific lipidomic and metabolic characterization at different time points during multi-day ex vivo perfusion of human kidneys. Through matrix-assisted laser desorption/ionization mass spectrometry imaging (MALDI-MSI) at subcellular spatial resolution with isotope tracing (39, 40), we show insights into the metabolic viability of discarded human kidneys following 4- and 8-day *ex vivo* preservation. Next, we investigated functional parameters of discarded human kidneys during 4-day ex vivo preservation and conclude by demonstrating that transplantation of a viable graft is technically feasible in a porcine auto-transplantation model.

In this proof-of-concept study we demonstrate that multi-day preservation of human kidneys is feasible at sub normothermia. Analysis of cell-type-specific metabolic dynamics during ex vivo kidney perfusion provided us with a more detailed understanding of the mechanisms underlying *ex vivo* perfusion.

## RESULTS

### 2.1 Sub normothermic organ culture platform

To preserve discarded kidneys for prolonged periods we developed an organ culture platform using porcine abattoir kidneys. The platform is built around a custom designed air-tight organ chamber (Figure 1A). Renal flow is provided by a pressure controlled centrifugal pump set at 75 mmHg. Arterial, venous and ureteral cannulation allowed continuous assessment of hemodynamic parameters and perfusate sampling. The kidneys are perfused with an acellular perfusate containing DMEM F12 and human serum albumin as main components. To support renal metabolism we utilized a perfusate rich in glutamine, citrate and acetate (for full composition see Table S1) (34, 37–39). Temperature was maintained at 25°C throughout the entire culture period. For oxygenation, the acellular perfusate was saturated with carbogen (95% O_2_, 5% CO_2_) through an oxygenator. Pilot experiments demonstrated that this method was proficient for oxygen delivery at sub normothermia (Figure S1).

**Figure 1.**
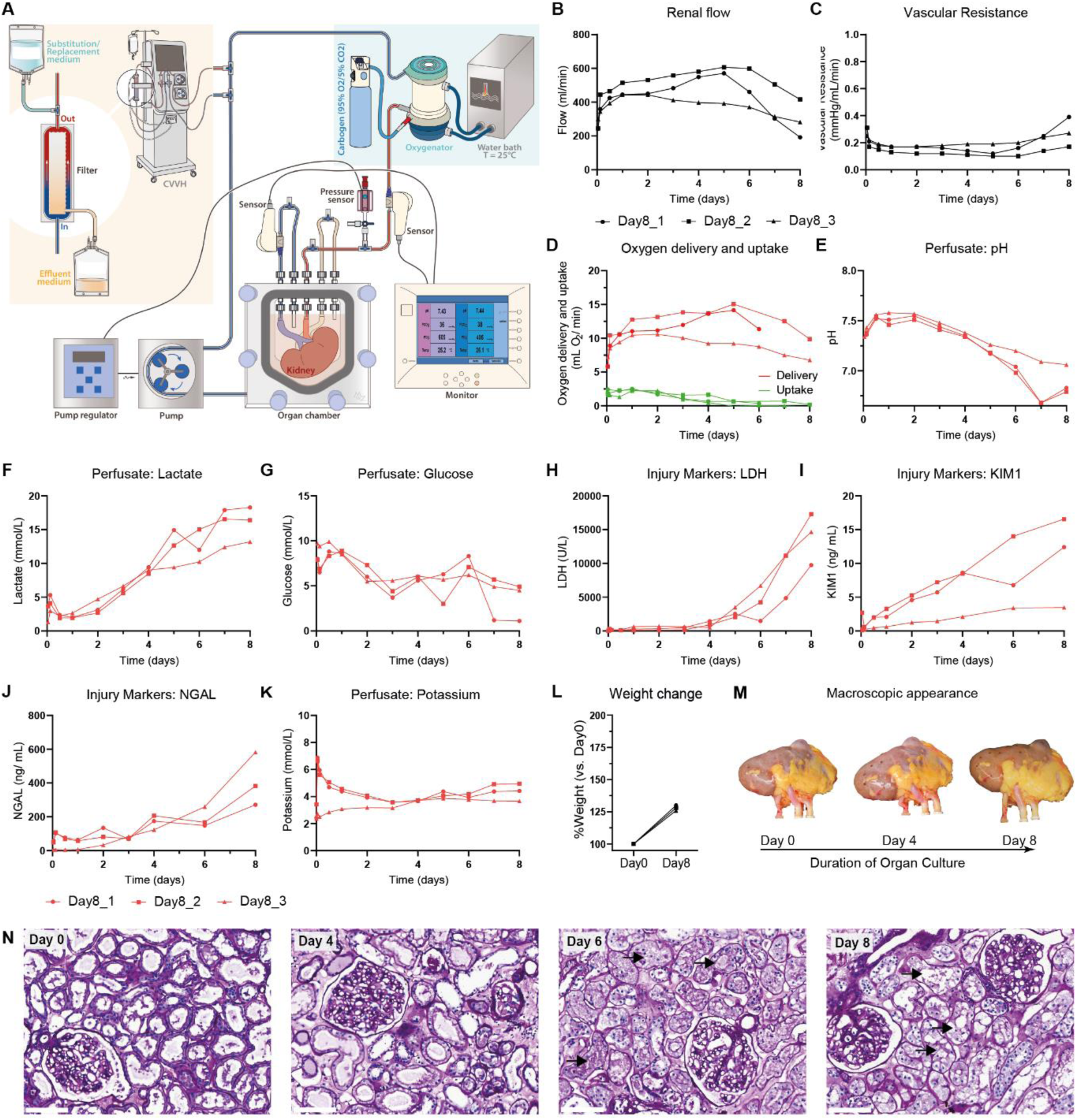
Perfusion dynamics during 8-day perfusion of human kidneys. Three discarded human kidneys were cultured for an 8-day period. **A**, Schematic overview of organ culture platform. Kidneys were preserved in a custom-designed organ chamber. A pressure controlled centrifugal pump set at 75 mmHg perfused the renal artery with acellular perfusate after it passed through an oxygenator. Perfusion temperature was maintained at sub normothermia (25°C) throughout the entire culture period. Arterial, venous and ureteral cannulation allowed continuous assessment of perfusion parameters. Urine was recirculated. Continuous hemofiltration allowed removal of small molecular weight waste products and substitution with fresh perfusate. **B-C**, Renal flow (**B**) and vascular resistance (**C**) throughout the 8-day period. **D-G**, Metabolic perfusate dynamics throughout the 8-day period. Oxygen delivery as calculated from the pO_2_ in the arterial inflow and oxygen uptake as calculated from the delta pO_2_ between the arterial inflow and venous outflow (**D**). Perfusate pH (**E**), perfusate lactate (**F**), and perfusate glucose (**G**) as measured in the arterial inflow. **H-K**, Perfusate injury markers throughout the 8-day period. Perfusate LDH (**H**) as markers for general cell damage. Perfusate KIM1 (**I**) and perfusate NGAL (**J**) as markers for proximal tubular and distal tubular injury, respectively. Because urine is recirculated tubular injury markers are measured within the perfusate (arterial inflow). Perfusate potassium as marker for general cell damage (**K**). **L**, Whole organ weight change (%) after the 8-day period as compared to baseline weight. **M**, Macroscopic appearance of discarded human kidneys throughout the 8-day period. One out of three representative kidneys shown. **N**, Representative images of PAS staining on cortex biopsies taken at different timepoints during the 8-day period (Day0, Day4, Day6, Day8). Black arrows highlight areas that demonstrate loss of tubular lumen. Bars represent 100 µm.

Urine was recirculated as it has previously been shown to improve hemodynamic control and electrolyte balance (14, 41, 42). To allow removal of metabolic waste products that otherwise accumulate in a closed system during prolonged culture, we incorporated continuous haemodialysis (Figure S2). Continuous haemodialysis was driven by an independent pump that allowed removal of small molecular waste products over a dialysis filter at a rate of 40 mL/hr. Substitution perfusate was added post-filter at the same rate (for composition see Table S1). This resulted in a continuous 5% fluid exchange. During multi-day pilot experiments with porcine kidneys, we observed that continuous haemodialysis improves perfusion hemodynamics (Figure S2).

### 2.2 Perfusion dynamics during 8-day culture

Next, we applied the platform to human kidneys that had been deemed unsuitable for transplantation (for donor data see Table S2). To test how long our platform could preserve kidneys, three discarded human kidneys were cultured for an extended period with real-time monitoring of multiple perfusion parameters. The perfusion was stopped after 8-days due to the observation of progressive disturbances in perfusion dynamics (Figure 1).

Upon connection of the kidneys to the culture platform, renal flow increased during the first hours of perfusion with vascular resistance decreasing (Figure 1B-C). Renal flow remained stable until Day5-Day6 of perfusion, thereafter demonstrating a gradual decrease. Oxygen delivery has a linear relationship with renal flow and remained between 5 and 15 mL O_2_/min throughout the 8-day period (Figure 1D). Oxygen uptake was constant during the first days and clearance of lactate was observed. After Day4, oxygen uptake decreased below 0.5 ml O_2_/ min (Figure 1D). In parallel, perfusate pH dropped below 7.30 for all three kidneys (Figure 1E) with a progressive increase in perfusate lactate towards Day8 (Figure 1F) whilst perfusate glucose remained within normal range (Figure 1G). Taken together this implies that the kidneys undergo a metabolic shift towards glycolysis.

Perfusate lactate dehydrogenase (LDH), an enzyme marker for cellular injury, revealed relatively low cellular injury until Day4 of perfusion, followed by a progressive increase that peaked on Day8 (Figure 1H). Tubular injury markers KIM1 and NGAL were measured within the perfusate given that urine was recirculated (Figure 1I-J). KIM1 showed a gradual increase throughout the 8-day period whereas NGAL showed the strongest increase between Day6 and Day8. Potassium concentration increased following the start of culture but remained within normal range thereafter for the 8-day period (Figure 1K). On Day8 kidney weight had increased with 28±1% compared to baseline (Figure 1L; Day0: 283±39, Day8: 363±52 gram). Absence of excessive oedema formation, as indicated by weight gain following perfusion, has previously been associated with preserved vascular barrier function (15). Macroscopic appearance at different timepoints throughout the 8-day period did not change (Figure 1M). Tissue histology on biopsies taken during perfusion demonstrated progressive damage to the apical membrane of tubular cells, including loss of tubular lumen between Day6 and Day8 (Figure 1N).

### 2.3 Lipidomics at single cell resolution highlight phenotypic remodelling

To gain insight into metabolic changes during sub normothermic organ culture, renal cortex punch biopsies were taken at five different timepoints throughout the 8-day period (Day0, Day2, Day4, Day6, Day8) and analysed through high spatial resolution matrix-assisted laser desorption/ionization mass spectrometry imaging (MALDI-MSI) as previously reported (39). This supports the simultaneous mapping of hundreds of lipids and metabolites per pixel while providing spatial information at subcellular resolution (5×5 µm^2^ pixel size, with an average renal cell diameter of approximately 10 µm) (43). Following MALDI-MSI measurements, we performed Uniform Manifold Approximation and Projection (UMAP) analysis based on the lipidomic data and post-MALDI-MSI immunofluorescence staining was conducted for cell type identification (Figure S3A). We could distinguish LTL+ proximal tubules, ECAD+ tubules and podocytes from other renal cell types based on specific lipid signatures (Figure 2A and Figure S3B), composed of phospholipids predominantly present in cell membranes. Proximal tubule, ECAD+ tubule and podocyte identity was confirmed by immunofluorescence staining on post-MALDI-MSI analysed tissue through alignment with cell type distribution (Figure S3A-B).

**Figure 2.**
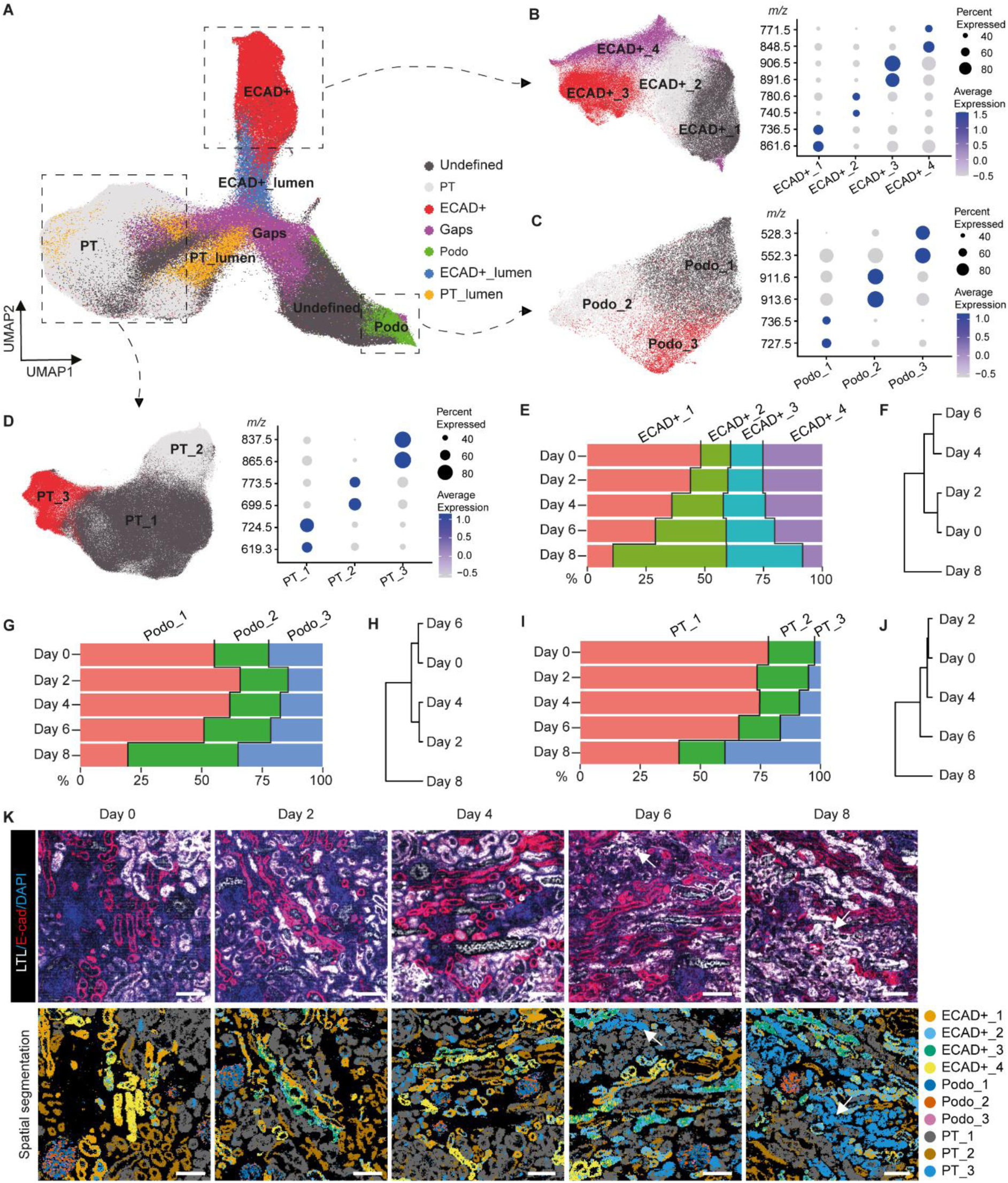
Spatial lipidomics reveals epithelial phenotypic remodeling during 8-day sub normothermic culture of human kidneys. **A**, Lipid heterogeneity in biopsies taken at different timepoints (Day0, Day2, Day4, Day6, Day8) from human kidneys (n=3) throughout the 8-day culture period allows identification of the main epithelial cell types, visualized in UMAP plot of MALDI-MSI data (5x5 µm^2^ pixel size). **B-D,** Sub clustering of ECAD+ tubular cells (**B**), podocytes (**C**), and proximal tubular cells (PT) (**D**) displays different epithelial phenotypes within each cell type. Dot plots display expression of cluster-specific lipid features. **E**, Percentage of each ECAD+ tubular phenotype in biopsies taken at different timepoints during 8-day culture. **F,** Hierarchical clustering of the percentage of ECAD+ tubular phenotypes in biopsies taken at different timepoints. **G**, Percentage of each podocyte phenotype in biopsies taken at different timepoints during 8-day culture. **H,** Hierarchical clustering of the percentage of podocyte phenotypes in biopsies taken at different timepoints. **I,** Percentage of each tubular PT phenotype in biopsies taken at different timepoints during 8-day culture. **J,** Hierarchical clustering of the percentage of PT tubular phenotypes in biopsies taken at different timepoints. **K,** Immunofluorescence staining on post-MALDI-MSI samples taken at different timepoints during 8-day organ culture (top panel). Spatial segmentation showing distribution of different renal epithelial phenotypes (bottom panel). White arrows highlight the area of PT_3. Bars represent 200 μm.

First, we evaluated the effect of 8-day ex vivo perfusion on cell phenotypic preservation by looking at changes in lipid species profiles. Unsupervised clustering was performed for the ECAD+ tubules (Figure 2B), podocytes (Figure 2C) and proximal tubules (PT) (Figure 1D), respectively. Different subsets of cell phenotypes were observed, characterized by different lipid signatures (Figure 2B-D). During 8-day culture a gradual change in relative composition of the different cell phenotypes was observed. Unsupervised hierarchical clustering of the different timepoints for each cell population demonstrates that lipid remodelling was predominantly observed between Day6 and Day8 of culture (Figure 2E,G,I). For example, phenotype PT_3, which was barely present on Day0, comprised >30% of proximal tubular cells on Day8 and was characterized by a lipid marker at *m/z* 865.6. This is an injured PT marker that we previously found following ischemia/reperfusion injury in a murine model (39). In parallel, a decrease of PT_1 and PT_2 phenotypes was observed from Day0 to Day8. Comparing the spatial segmentation with immunofluorescence staining revealed that this abnormal PT_3 phenotype mainly colocalizes with LTL staining in Day6 and Day8 biopsies (Figure 2K), confirming the phenotypic changes within proximal tubules. These observations suggest that cellular phenotypes are preserved up to Day4-Day6 of ex vivo preservation.

To further assess how the relative abundances of lipids change during the 8-day culture, we compared the lipid species characteristic for the main tubular subsets (Figure 3A-B). Due to the variation between different donors, average lipid peak intensities were normalized to their level at Day0. Spatial distributions and relative abundances of the various lipid markers for the main renal cell types remained similar until Day6 of culture (Figure 3A). This suggests preservation of the main cell membrane lipid components without degradation up to Day6-Day8 of ex vivo perfusion.

**Figure 3.**
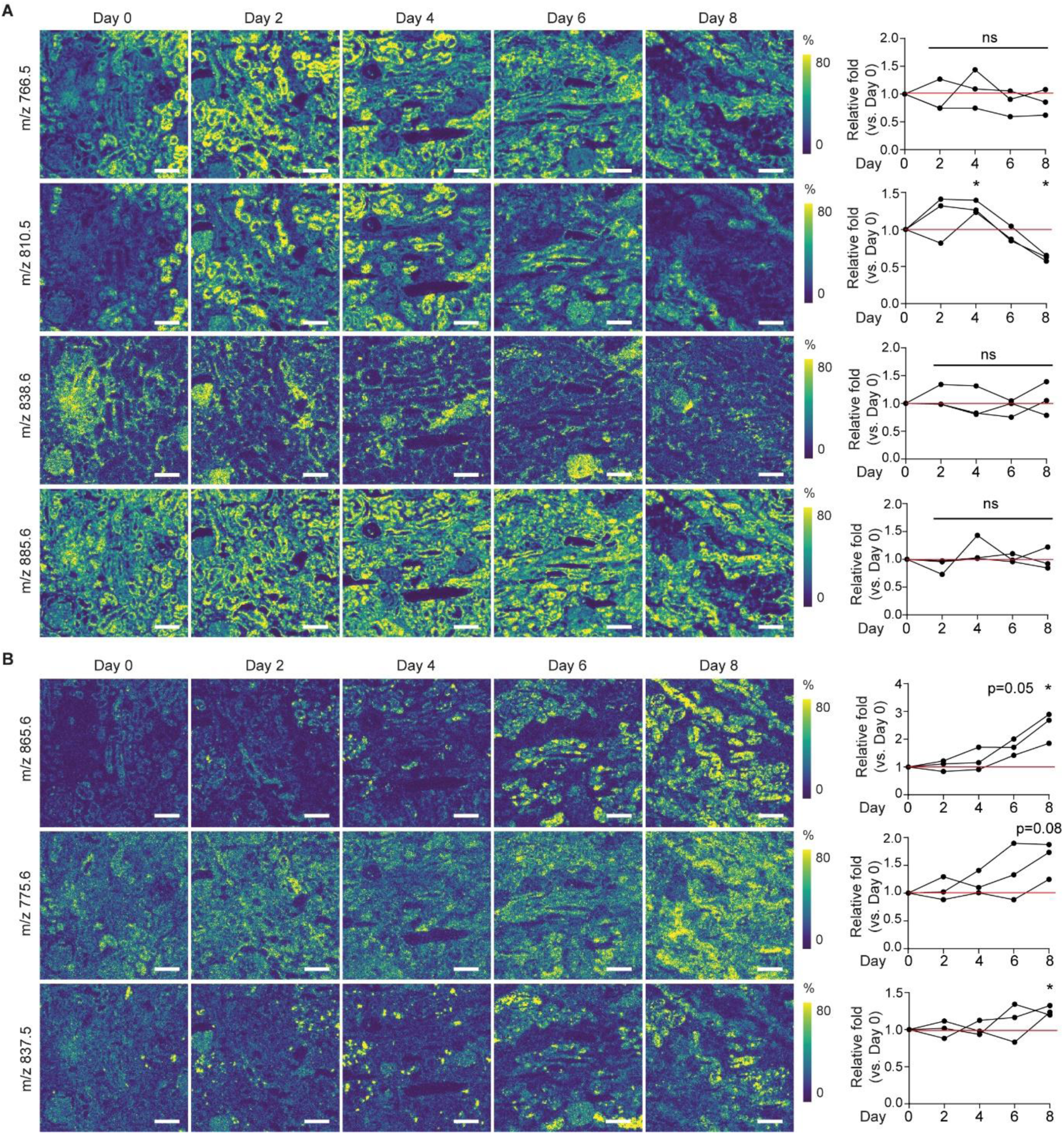
Spatial lipid species distribution changes during 8-day sub normothermic culture of human kidneys. **A**, Representative images demonstrate changes in spatial distribution of lipid species characteristic for normal renal cell phenotypes at different timepoints during 8-day organ culture, as recorded by MALDI-MSI (5×5 µm^2^ pixel size). Lipid species *m/z* values are representative for PT_1 and PT_2 (*m/z* 766.5), PT_1 (*m/z* 810.5), podocytes (*m/z* 838.6), and all tubular structures (*m/z* 885.6). Right panel shows relative fold change compared to baseline (Day0) for each timepoint during culture. Scale bar = 200 μm. **B**, Representative images demonstrate changes in spatial distribution of lipid species that are characteristic for abnormal PT phenotype (PT_3) at different timepoints during 8-day organ culture, as recorded by MALDI-MSI (5x5 µm^2^ pixel size). Right panel shows relative fold change compared to baseline (Day0) for each timepoint during culture. Bars represent 200 μm. *p < 0.05. one sample T test. All images generated from same samples as shown in Figure 3K with IF staining, spatial segmentation. Bars represent 200 μm.

On the contrary, the relative abundance of lipid markers characteristic for the PT_3 proximal tubules was significantly higher on Day8 as compared to Day0 (Figure 3B). Visualization of their spatial distribution and changes in their relative abundance revealed that this lipid remodelling started between Day4 and Day6 and progressively worsened, as is illustrated by a more homogenous distribution throughout the tissue towards Day8.

To gain insight in what underlies this change in lipid species profile we assessed lipid peroxidation. A progressive increase in perfusate malondialdehyde (MDA) was observed after Day6 of perfusion (Figure 4A), indicating progressive oxidative stress. The increased abundance of oxidized lipids in the perfusate on Day8 as compared to Day0 till Day4 was confirmed through targeted lipidomics (Figure S4). Whereas the abundance of oxidized lipids within the perfusate was low until Day4-Day6 of perfusion, a considerable increase was observed on Day8 of perfusion. Next, untargeted lipidomics was performed on 8-day perfused kidney tissue and compared with unperfused control samples. In Day8 tissue, the lipids demonstrating the most significant fold change are all oxidized phospholipid species (Figure 4B). Spatial distribution of the oxidized phospholipids was assessed within the MALDI-MSI analysed biopsies (Figure 4C). This supports that oxidized lipid species are absent between Day0 and Day4-Day6 of ex vivo perfusion, with progressive oxidative stress taking place between Day4-Day6 and Day8 of perfusion. Moreover, comparing the spatial distribution of the oxidized phospholipid PE(O-36:2) on Day8 of perfusion (Figure 4C) with the spatial segmentation of the different cell clusters (Figure 2K) demonstrates that it co-localizes with the PT_3 proximal tubules.

**Figure 4.**
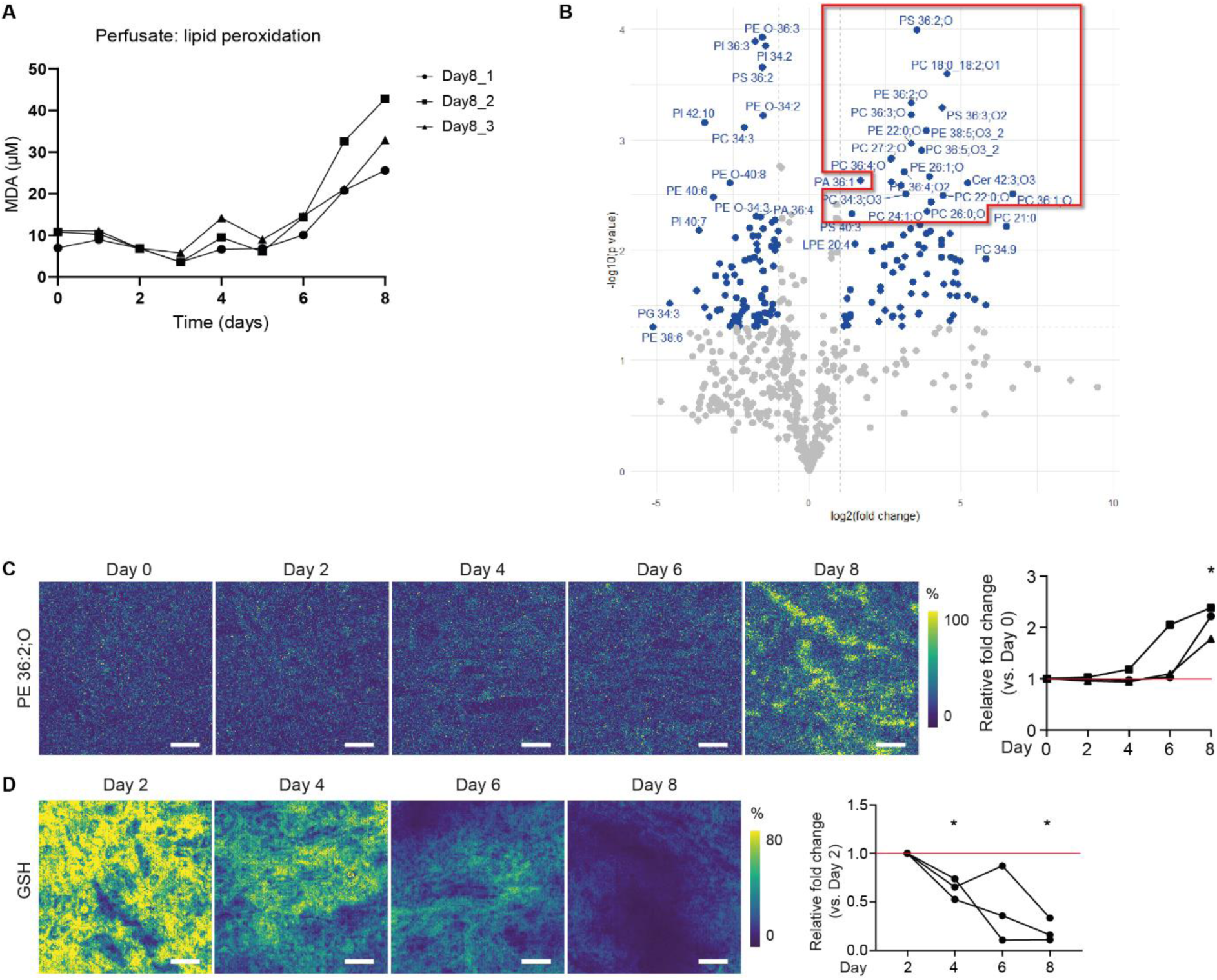
Oxidative damage during 8-day perfusion. **A**, Perfusate malondialdehyde (MDA) as marker for lipid peroxidation during 8-day perfusion. **B**, Volcano plot comparing fold change in all lipid species measured in 8-day perfused tissue samples with control kidney tissue (n=3 human kidneys per group), as identified through untargeted lipidomics. **C**, Representative images demonstrate changes in spatial distribution of oxidized phospholipid species (PE 36:2;O) at different timepoints during 8-day perfusion, as recorded by MALDI-MSI (5x5 µm^2^ pixel size). Right panel shows relative fold change compared to Day0. *p < 0.05. One sample T test. **D**, Representative images for changes in the anti-oxidant glutathione (GSH) at different timepoints during 8-day perfusion. Right panel shows relative fold change compared to Day2. *p < 0.05. One sample T test.

The observed increase in oxidative stress is supported by the reduced relative abundance of the anti-oxidant glutathione (Figure 4D). Due to the variation of metabolite level likely caused by varying cold ischemic periods before commencing perfusion, we normalized the reduced glutathione average peak intensity to its level at Day2 to compare its relative abundance during prolonged perfusion. A gradual decrease in glutathione abundance was observed during the period of ex vivo perfusion (Figure 4D). Together with the decreased oxygen consumption (Figure 1D) and increase in oxidized lipid species (Figure 4, Figure S4), this implies that oxidative stress caused by mitochondrial uncoupling, progressively worsens after Day4 of perfusion. Therefore, we next assessed metabolic activity after 4-and 8-day ex vivo perfusion using spatially resolved isotope tracing.

### 2.4 Isotope tracing indicates preserved metabolic activity during prolonged perfusion

Defective metabolism and metabolic recovery play a key role in the response to kidney injury (35, 36). To assess preservation of central carbon metabolism and changes in nutrient partitioning during 8-day culture we applied our recently described spatial dynamic metabolomics platform (39). In short, dedicated cortex biopsies were taken on Day0, Day4 and Day8 from the three human kidneys during 8-day perfusion. These biopsies were incubated with ^13^C-labeled glucose or glutamine for 2 h at 37°C. MALDI-MSI was then used to spatially visualize ^13^C enrichment of downstream intermediates of glycolysis and the TCA cycle at subcellular resolution.

We found that after 4- and 8-days of ex vivo perfusion all renal cell types identified in the biopsies maintain active cell metabolism and can use glucose and glutamine as nutrient sources for glycolysis and the TCA cycle (Figure 5). The enrichment of the ^13^C_6_-glucose-derived isotopologues, ^13^C_3_-3PG, ^13^C_3_-Lactate and ^13^C_2_-Glutamate demonstrated active glycolysis and contribution of glucose to the TCA cycle, respectively (Figure 5A-F). The enrichment of ^13^C_5_-glutamine-derived isotopologues, ^13^C_5_-Glutamate, ^13^C_4_-Succinate, ^13^C_4_-Malate and ^13^C_3_-Glutamate, demonstrated passage through the entire oxidative TCA cycle (Figure 5H-N). A partial contribution of glutamine to reductive TCA cannot be ruled out.

**Figure 5.**
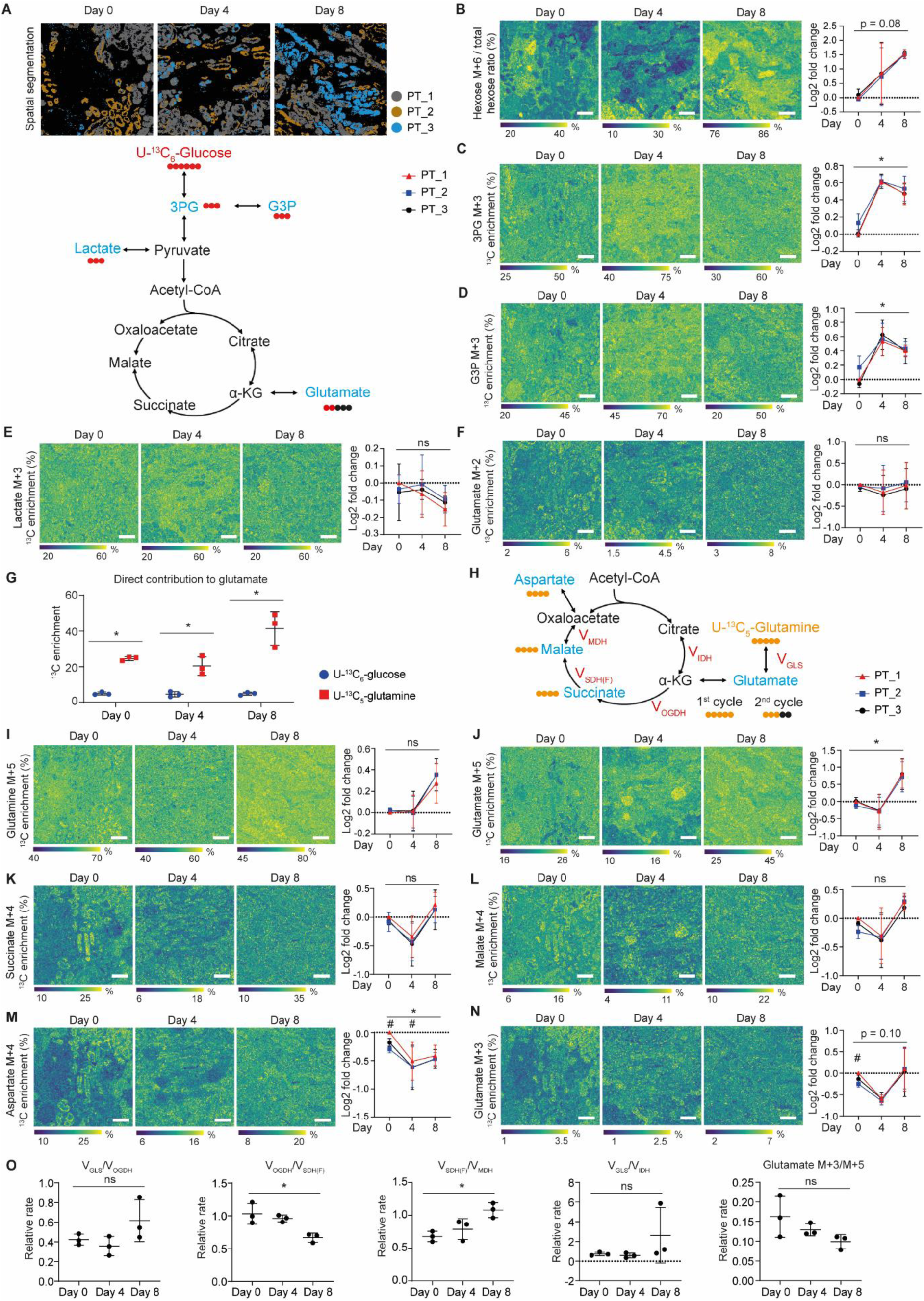
Spatial dynamic metabolic measurements on biopsies taken at different timepoints during 8-day organ culture. **A**, Spatial segmentation showing distribution of different proximal tubular phenotypes. **B-F,** Images and graphs showing the spatial dynamic metabolic measurements using U-^13^C_6_-glucose on the biopsies obtained at different timepoints during the 8-day organ culture period after 2 hours’ incubation. Graphs are shown as log2 fold change compared to PT1 at Day 0. Bars represent 200 μm. *p < 0.05. One way ANOVA test. **G,** Direct carbon contribution of different nutrients to glutamate in proximal tubule, as measured from the glutamate isotopologues M+2, M+3, and M+5. *p < 0.05. two-tailed unpaired t test. **H-N,** Images and graphs showing the spatial dynamic metabolic measurements using U-^13^C_5_-glutamine on the biopsies obtained at different timepoints during the 8-day organ culture period after 2 hours’ incubation. Graphs are shown as log2 fold change compared to PT1 at Day 0. Bars represent 200 μm. *p < 0.05. One way ANOVA test. #p < 0.05. two-tailed paired t test compared to PT1. **O**, Relative flux rate of different TCA steps. *p < 0.05. One way ANOVA test. Data are represented as mean±SD. All images generated from same samples as shown in Figure 3K with IF staining, molecular histology and spatial segmentation.

Subsequently, we focused on metabolic changes in the different proximal tubule phenotypes identified during 8-day culture (Figure 5A). First, analysis of the isotopologues derived from ^13^C_6_-glucose showed all PT phenotypes after perfusion to have gradually higher U-^13^C_6_-hexose and higher ^13^C_3_-3PG enrichment compared to PT at Day0 (Figure 5B-C). A higher ^13^C_3_-G3P enrichment was also observed after perfusion, pointing to a side branch of glycolysis (Figure 5D). The enrichment of ^13^C_3_-lactate and ^13^C_2_-glutamate from glucose carbons show no differences (Figure 5E-F). These data suggest an active glycolysis pathway and the contribution of glucose to the TCA cycle until Day8 of perfusion. The carbon contribution of ^13^C_5_-glutamine to glutamate is about 5 times higher than ^13^C_6_-glucose (Figure 5G). As the conversion of glutamate to α-ketoglutarate (α-KG) is very significant in mitochondria, this shows a higher contribution of glutamine to the TCA cycle than glucose. The detection of ^13^C-labeled TCA cycle intermediates derived from ^13^C_5_-glutamine demonstrates that the TCA cycle remained active upon till Day8 of perfusion (Figure 5H-N). At Day0 PT_1 and PT_2 (PT_3 barely exists at Day0) displayed heterogeneity in their metabolic dynamics, as shown by ^13^C_5_-glutamine-derived ^13^C_4_-aspartate and ^13^C_3_-Glutamate (Figure 5M-N). However, this heterogeneity between different PT phenotypes was lost during perfusion, especially at Day8 (Figure 5M-N). There was a significant increase in ^13^C_5_-glutamate and decrease in ^13^C_4_-aspartate in PT at Day8 as compared to Day0 (Figure 5J, 5M). To assess the relative enzyme activity in the TCA cycle at different timepoints of perfusion, we performed Q-Flux analysis (^13^C_5_-glutamine-based method) using equations from a recent publication where this method was validated (44). This analysis demonstrated that the fractional contribution of succinate-to-succinate dehydrogenase forward flux was decreased at Day8, while the succinate dehydrogenase forward flux relative to malate dehydrogenase flux was increased at Day8 (Figure 5O). Overall, there is a decreasing trend in the fraction of ^13^C_3_-glutamate that is derived from ^13^C_5_-glutamate through the entire oxidative TCA cycle (Figure 5O).

Most importantly, we observed no differences in the relative rate of TCA cycle enzymes between Day0 and Day4 of perfusion (Figure 5O). This underpins that sub normothermic machine perfusion allows multi-day ex vivo metabolic preservation of diseased human donor kidneys.

### 2.5 Renal function during 4-day culture

Following the demonstration of metabolic preservation for more than 4-days during sub normothermic perfusion we included an additional five discarded human kidneys (for donor data see Table S2). These kidneys were cultured for a 4-day period during which perfusion dynamics, renal function and tissue integrity were assessed (Figure 6). Renal flow and vascular resistance were stable throughout the 4-day period (Figure 6A-B). Oxygen uptake was maintained around 1 mL O_2_/min (Figure 6C-D). pH was within normal range after 3 h of perfusion, and ended at 7.37±0.05 on Day4 of perfusion (Figure 6E). In contrast to the accumulation of injury markers observed during the 8-day perfusion (Figure 2), only a minor increase in perfusate LDH, KIM1 and NGAL was observed throughout the 4-day period (Figure 6H-J). Histological scoring of periodic acid-Schiff (PAS) staining was performed for the five discarded human kidneys that were perfused for the 4-day period (Figure 6K, Figure S5). To this end wedge biopsies were taken at the end of 4-day perfusion and control biopsies from the contralateral kidneys which we received for Day4_1 and Day4_2.

**Figure 6.**
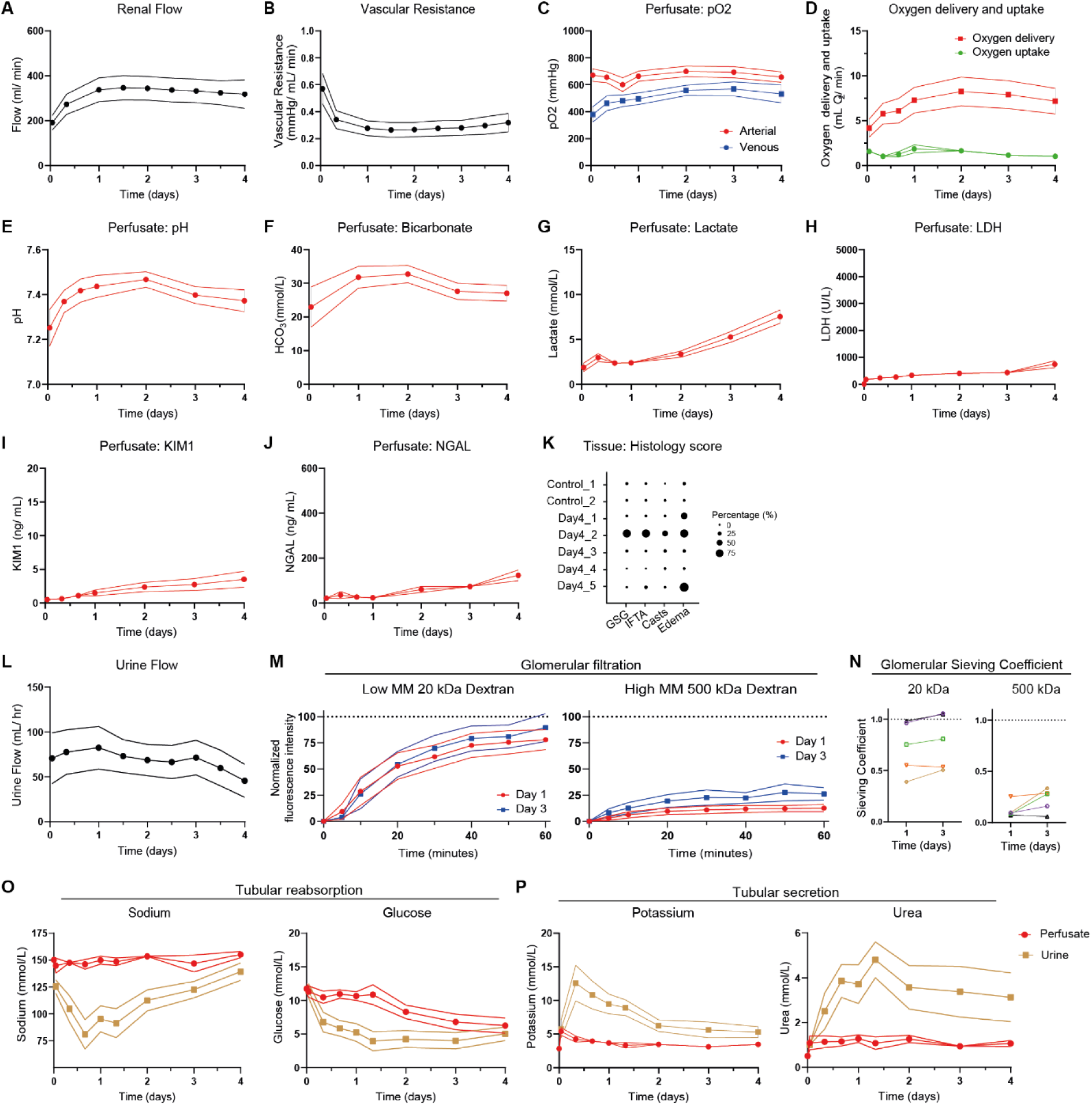
Perfusion dynamics and function during 4-day sub normothermic culture of eight human kidneys. **A-B**, Course of renal flow (**A**) and vascular resistance (**B**) during the 4-day perfusion period. **C-G**, Metabolic perfusate dynamics. Perfusate pO_2_ (**C**) measured in the arterial inflow and venous outflow. Oxygen delivery (**D**) as calculated from the pO_2_ in the arterial inflow and oxygen uptake (**D**) as calculated from the difference in pO_2_ between the arterial inflow and venous outflow. Perfusate pH (**E**), perfusate bicarbonate (**F**), and perfusate lactate (**G**) as measured in the arterial inflow. **H-J**, Perfusate injury markers during the 4-day perfusion period. Perfusate LDH (**H**) as marker for general cell damage. Perfusate KIM1 (**I**) as marker for proximal tubular cell damage. Perfusate NGAL (**J**) as marker for distal tubular cell damage. **K**, Tissue histology scoring of the five discarded human kidneys that were cultured for a 4-day period. Day4_1 and Day4_2 are the contralateral kidneys of Control_1 and Control_2, respectively. Representative PAS staining’s can be found in Figure S5. **L-P**, Renal function during 4-day perfusion. **L**, Urine flow. **M**, Assessment of glomerular filtration barrier on Day1 and Day3 of perfusion. Low molecular mass 20 kDa fluorescent dextran (FITC-labelled) and high molecular mass 500 kDa fluorescent dextran (TRITC-labeled) were infused with subsequent perfusate and urine collection (n=5 human kidneys). **N**, Glomerular sieving coefficient for 20 kDa and 500 kDa dextran (n=5 human kidneys). For **M** and **N**, fluorescence intensity of dextran in the urine was normalized for fluorescence intensity of dextran in the perfusate. **O-P**, Perfusate and urine concentration differences demonstrates tubular reabsorption of sodium and glucose (**O**) and secretion of potassium and urea (**P**) during perfusion at sub normothermia (25°C). Data are presented as mean±SEM.

Finally, we assessed renal function during ex vivo preservation. In all kidneys glomerular filtration was observed, as indicated by urine production (Figure 6I). To assess the integrity of the glomerular filtration barrier fluorescently labelled dextran’s of 20 kDa and 500 kDa were added to the perfusate on Day1 and Day3 of culture. The 20 kDa dextrans were filtered into the urine whereas the 500 kDa dextrans were retained within the vascular compartment, indicating preserved barrier function (Figure 6J-K). We observed active tubular transport during sub normothermic culture, as indicated by reabsorption and secretion of electrolytes and metabolites (Figure 6L-M). Sodium and glucose were reabsorbed (Figure 6L) whereas potassium and urea were secreted by the kidneys (Figure 6M) during 4-day sub normothermic preservation. During the 8-day perfusions, kidneys lost their ability to create electrolyte gradients between Day4 and Day6 of preservation (Figure S6).

### 2.6 Early phase in vivo reperfusion

To demonstrate that it is feasible to procure, preserve for 4-days and transplant a kidney, we performed a porcine auto transplantation as proof-of-concept (Figure S7, n=1). Following 4-day ex vivo perfusion the kidney was auto-transplanted into the right groin of the same pig. The renal vein was anastomosed end-to-side to the caval vein and the renal artery was anastomosed end-to-side on the distal aorta. During a 3 hour observation period, macroscopic appearance of the kidney continued to show a pink, well vascularized organ. Vascular anastomosis patency was confirmed through doppler ultrasound with visible flow through the anastomoses and within the kidney (Figure S7A). The ureter of the auto-transplanted kidney was cannulated and selective collection demonstrated that 8 mL of urine was produced by the transplanted graft during this early phase of reperfusion.

## DISCUSSION

In this study, we developed a kidney culture platform that combines perfusion at sub normothermia (25°C) with an acellular perfusate rich in metabolites that serve as direct TCA cycle intermediates, urine recirculation, and continuous hemofiltration. Following an initial phase of optimalization using porcine kidneys, eight discarded human kidneys were cultured up to 8-days. Using spatially resolved single cell resolution isotope tracing we demonstrate active metabolism in all the different renal cell types over this period. Beyond Day4-Day6 of perfusion, composition of nephron segments, as assessed through spatial lipidomics, demonstrated substantial changes in lipid composition and considerable increases in perfusate injury marker LDH were observed. Up to 4-days perfused human kidneys, however, maintained functional parameters.

Characterization of metabolic and lipidomic changes during multi-day perfusion was performed through MALDI MSI, allowing the simultaneous mapping of hundreds of lipids and metabolites per pixel while providing spatial information at subcellular resolution (5×5 µm^2^ pixel size). Combining this with isotope tracing allowed us to observe cell-type-specific metabolic changes during multi-day perfusion and correlate this with whole organ perfusion parameters. (40, 43). Spatial lipidomics allowed the identification of different cell type subpopulations, and their relative changes in %composition during 8-day perfusion. Within the cluster of LTL-positive PTs, PT_3 was barely present on Day0 whereas highly abundant on Day8. Isotope tracing was performed in vitro at 37°C, thereby simulating a 2h rewarming period. Cell-type-specific metabolic changes were assessed based upon ^13^C_6_-glucose and ^13^C_5_-Glutamine nutrient partitioning. This yielded unexpected insights into the metabolic activity at different timepoints following prolonged *ex vivo* preservation. We initially expected to observe a decrease in %-enrichment of TCA cycle metabolites on Day8 of perfusion as compared to Day0 and Day4, because of loss of cell viability. This was expected from the accumulation of cell injury (Figure 2G-I), decrease in oxygen uptake between Day4 and Day8 (Figure 2D), increased lactate abundance (Figure 2E), and decrease in perfusate pH (Figure 2D). However, ^13^C-isotope tracing during *in vitro* rewarming revealed persistent metabolic activity up to Day8 of perfusion, in both glycolysis and TCA cycle (Figure 5A and 5C). However, the initial spatial heterogeneity in TCA metabolism in the various tubular epithelial segments was lost after 4 days and some reductive TCA activity can also not be excluded. We therefore also performed Q-flux analysis, a recently described method to assess mitochondrial metabolism (44), which demonstrated no differences in the relative rate of TCA cycle enzymes between Day0 and Day4 of perfusion (Figure 6O).

All kidneys maintained normal functional parameters up to 4-days of perfusion. The glomerular filtration barrier was assessed through the infusion of small (10 kDa) and large (500 kDa) molecular weight (MW) labelled Dextrans. Large MW dextrans were retained within the vascular compartment, whereas the small MW dextrans were filtered into the urine. Active tubular transport machinery throughout the 4-day period was confirmed by electrolyte gradients between perfusate and urine. Sodium and glucose were reabsorbed, whereas potassium and urea were excreted. Renal reabsorption and secretion was comparable if not more pronounced during sub normothermic preservation as compared to a recent case report describing 48 hours normothermic human kidney preservation in which gradients were visualized the same way (17). Our data demonstrating active metabolism and functionality up to 4 days after procurement are also remarkable from the perspective that these were discarded human kidneys that had pre-existing damage and injury. For example, one of the kidneys was a retransplant organ.

Two decades ago, Brasile et al. was the first to demonstrate that it is feasible to preserve canine and human kidneys up to 48h within a sub normothermic (32°C) perfusion platform (15, 27, 45). They postulated that warm perfusion holds the potential to ameliorate reperfusion injury and recover function ex vivo (45), whereas cold preservation hampers (metabolic) recovery and results in cold-induced cellular injury (46, 47). When canine kidneys were warm perfused for 48 h at sub normothermia and reimplanted, immediate function was observed (15). Subsequently, the effect of renal function after severe warm ischemia (120 minutes) was assessed. Primary non-function (PNF) was observed for all control kidneys, that were either immediately transplanted or were 18 h cold preserved (HMP). Contrarily, 18h sub normothermic perfused kidneys reperfused well and functioned within minutes of reperfusion (27). Warm sub normothermic perfusion also improved function upon reimplantation as compared to cold preservation only (45). Therewith, cold ischemia was demonstrated as limiting factor, and sub normothermic perfusion as potential solution, for expanding the donor organ pool by including (warm ischemia) damaged kidneys. However, this work received little follow up by other groups since then. Our data corroborate in human kidneys their initial observations and show that using the sub normothermic approach in combination with our optimized organ perfusion platform one can extend organ preservation into days.

Nowadays, most-if not all,-groups in the field of organ preservation focus upon preservation of the kidney at body temperature. Reports of successful perfusion up to or beyond 24h are rare (9, 14, 17). We postulate that this is due to high (and heterogenous) metabolic preferences of the kidney at body temperature and nephrotoxic effects due to free haemoglobin accumulation following haemolysis when using oxygen carriers. Indeed, Hosgood et al. recently demonstrated that free heme levels significantly increased after only 1 h of perfusion (48), being (just) below potentially nephrotoxic levels (20, 21, 49).

In sum, we demonstrate that perfusion at sub normothermia supports prolonged ex vivo preservation, thereby creating a platform for (complex) therapeutic and regenerative interventions in metabolically active kidneys. The ability to preserve kidneys in a metabolically active state for days instead of hours may allow the implementation of technologies that can be game-changers for transplantation. Donor organs are already routed through specialized facilities (Organ Perfusion and Regeneration (OPR)-units) for organ assessment and if necessary pre-treatment by means of relatively short periods of machine perfusion (5). Breakthroughs in liver preservation are demonstrating the range of possibilities for prolonged graft preservation (6, 7, 13), including transplantation of livers that would otherwise have been discarded (50), transplantation after 3-day ex vivo preservation (12), immuno-modulation during machine perfusion (13), and regeneration of bile ducts following cholangiocyte organoid transplantation (51).

## MATERIALS AND METHODS

### Study design

The primary objective of this study was de develop an organ culture platform that would support multi-day ex vivo preservation of metabolically active human kidneys. The study was carried out in three phases. First, we optimized our platform using porcine kidneys from a local abattoir (see supplementary materials and methods). We found that sub normothermic preservation at 25°C in a custom-made platform that includes acellular perfusate enriched with TCA cycle intermediates, urine recirculation, and continuous hemofiltration allows multi-day preservation of porcine kidneys. Next, three discarded human kidneys were cultured within the platform to determine for how many days it would support ex vivo preservation. All three human kidneys were stopped after an 8-day period due to progressive disturbances in perfusion dynamics. As detailed below, we applied our recently described single-cell-level spatial dynamic metabolomics platform on biopsies taken at different time points during 8-day culture to assess renal metabolic preservation. This allowed combined assessment at the whole-organ-level through perfusion dynamics and single-cell-level assessment to determine the effect of multi-day culture and cellular phenotype and metabolic viability. Finally, an additional set of five human kidneys was cultured for 4-day period, as was determined to be possible in the second stage.

### Human kidneys

A total of eight human kidneys were obtained for organ culture using the described platform, after being declined for transplantation because of various reasons (Table S1 for donor data). Research consent was given for all human kidneys before organ retrieval. No further research and ethics approval was required. The included kidneys were procured from deceased donors according to the Dutch national guidelines. After in situ flushing of the abdominal organs with cold University of Wisconsin (Belzer UW) preservation solution, the kidneys were retrieved and transported to the Leiden University Medical Center on Static Cold Storage (SCS) or Hypothermic Machine Perfusion.

Upon arrival at the laboratory, the renal artery, vein and ureter were cannulated using Luer lock connectors (*Cole Parmer, Barendrecht, the Netherlands*) whilst the organ culture platform was setup in parallel. Excessive renal fat was removed if necessary. Shortly before the start of culture kidneys were flushed with approximately 250 mL of cold DMEM F12 to remove the preservation solution and globally disinfected with betadine. Within the culture platform the renal artery, vein and ureter were connected to their designated outlets.

### Organ Culture Platform

The platform consists of a closed-loop circuit connected to a custom-designed air tight organ chamber (Mascal Design) that also serves as reservoir. Hemodynamic control is based on arterial pressure. A centrifugal pump (Masterflex L/S Digital Drive 600 rpm) perfused the renal artery through silicone tubing (LS25, Masterflex Metrohm) at a mean arterial pressure (MAP) of 75 mmHg. Pressure in the renal artery was measured in-line with a single-use pressure sensor (Edwards Lifesciences). The perfusate was oxygenated with a carbogen mixture of 95% O2 and 5% CO2 through a membrane oxygenator (Maquet Quadrox-I neonatal, Gettinge; or Sorin Lilliput 2, LivaNova) that was connected to a water bath set at set at 25°C for temperature regulation.

With urine being recirculated a Prismaflex system (Baxter) was connected in parallel to remove metabolic waste products and supply fresh nutrients whilst maintaining electrolyte homeostasis. A pediatric filter (Prismaflex HF20 set, Baxter) allowed the exchange of small molecular weight molecules. Blood flow through the filter was set at 20 mL/min. Fluid and small molecular weight molecules were removed at 40 mL/h over the filter. Fresh perfusate was substituted post-filter at the same rate. Thus continuous hemofiltration was performed at a filtration fraction of 5% (52).

Two in-line blood-gas sensor (CDI 500 system, Terumo Cardiovascular Systems) allowed continuous monitoring of pO_2_, pCO_2_, pH and temperature in the arterial inflow and venous outflow. Perfusate sampling ports were connected through a 3-way tap (Discofix, Braun) present on the arterial-, venous, and ureteral in-and outlet.

### Perfusate components

A detailed description of the perfusate composition is provided in Table S2. Organ culture was commenced with a total perfusate volume of approximately 700 mL. In brief, DMEM/F-12 (Gibco) was supplemented with Human Serum Albumin (Alburex 20, CSL Behring bv), Insulin-Transferrin-Sodium Selenite (ITS 100x, Sigma-Aldrich), sodium bicarbonate (Gibco), citric acid (Calbiochem, Merck), acetic acid (EMSURE, Merck), penicillin-streptomycin (Gibco), ciprofloxacin (Fresenius Kabi) and fungizone (Bristol-Myers Squibb). pH was adjusted with sodium hydroxide (Calbiochem, Merck) until within range (Target 7.30-7.45). Sodium levels were adjusted with distilled water (Sterile water, Versylene Fresenius) until within range (Target: 130-145 mmol L^-1^).

Continuous hemofiltration was performed at 40 mL/h. Substitution solution was added post-hemofilter by the Prismaflex system. The substitution solution contained the same components as the culture perfusate except for Human Serum Albumin (for composition see Table S1). Additional glucose (1M solution; D-Glucose, G8270, Sigma) was supplemented when perfusate glucose levels dropped below 4 mmol L^-1^, with the goal of maintaining euglycemia.

### Perfusate sampling and measurements

Perfusate sampling was performed at least three times per day throughout the culture period. Arterial, venous and urine samples were measured directly for Blood-gas analyses and monitoring of electrolytes and metabolites. Samples were aliquoted into Falcon tubes (1 mL) and frozen at -20°C and/or -80°C for later analysis.

An iSTAT1 blood analyzer (Abbott) was used for point-of-care blood-gas analysis (pH, pO_2_, pCO_2_, HCO_3_^-^). Perfusate samples from the arterial inflow and venous outflow were immediately measured using CG4^+^ iSTAT cartridges (Abbott). Blood-gas analysis data were used to calibrate the in-line shunt sensors (CDI-500 system, Terumo Cardiovascular Systems). Oxygen delivery (mL O_2_ min^-1^) was calculated as Arterial pO_2_ (mmHg) x Solubility of O_2_ (0.0031 mL O_2_ dL fluid^-1^ mmHg^-1^) x Renal Flow (dL min^-1^). Oxygen uptake (mL O_2_ min^-1^) was calculated as ΔpO_2_ (mmHg) x Solubility of O_2_ (0.0031 mL O_2_ dL fluid^-1^ mmHg^-1^) x Renal Flow (dL min^-1^). CG4+-measured pO_2_ values were used for calculation of oxygen delivery and uptake. Perfusate and urine electrolyte (Sodium, Potassium, Chloride) and metabolite (Glucose, Lactate, Urea) concentration were measured using Chem8 iSTAT cartridges (Abbott). The Clinical Chemistry Laboratory within the LUMC (re)measured sodium, potassium, urea and glucose within arterial perfusate and urine samples according to standard operating procedures, that had previously been stored at -80°C. Lactate Dehydrogenase (LDH) concentration was measured in perfusate samples.

Neutrophil gelatinase-associated lipocalin (NGAL) and kidney injury molecule-1 (KIM1) levels in the perfusate were measured using a quantitative sandwich enzyme immunoassay technique with NGAL Quantikine ELISA kit (DLCN20, R&D systems) and KIM-1 DuoSet ELISA kit (DY1750B, R&D systems) according to manufacturer’s instruction in samples that had previously been frozen and stored at -80°C. A thiobarbituric acid reactive substances (TBARS) assay kit (CaymanChemical, 10009055) was used to measure levels of malondialdehyde (MDA) in the perfusate according to manufacturer’s instructions in samples that had previously been frozen and stored at -80°C.

### Glomerular sieving coefficient

Filtration of high (500 kDa) and low (20 kDa) molecular mass dextran molecules was assessed to evaluate the intactness of the glomerular basement membrane during ex vivo preservation in the five human kidneys that were cultured for four days (53).

500 kDa TRITC labelled dextran (0.1 mg/ml) (Sigma-Aldrich 52194) and 20 kDA FITC labelled dextran (0,01 mg/ml) (Sigma-Aldrich 95648) were added to the perfusate after 24- and 72 hours of culture. Arterial and urine perfusate samples were taken every 10 minutes after dextran infusion for 1 hour. The fluorescence signal of the arterial and urine perfusate samples was directly measured using a Spectramax M5 (Molecular devices) at wavelengths 495-525 and 555-575.

### Tissue sampling and processing

Tissue punch biopsies (4mm) were taken at different time points during 8-day organ culture (Day0, Day2, Day4, Day6, Day8). At each timepoint biopsies were cut longitudinally into two pieces of which one was fixed in 4% paraformaldehyde and the other snap frozen in liquid nitrogen and stored at -80°C for further analysis. On Day0, Day4 and Day8 additional biopsies were taken for dynamic metabolic measurements. Tissue biopsies were placed into culture plates and incubated in a well-defined medium (Glucose free and glutamine free DMEM medium (Gibco, A1443001)), supplemented with 2% FCS, 5 mM glucose, 500 µM glutamine and penicillin/streptomycin (pH adjusted to 7.4) for 2 hours at 37°C and 5% CO_2_. For the ^13^C-labeling incubation, same amounts of either U-^13^C_6_-glucose (99%, Sigma, 389374) or U-^13^C_6_-glutamine (99%, Cambridge Isotope Laboratories, Inc. CLM-1822-H) were used to replace similar un-labeled nutrients in each medium. In the end, tissue slices were quenched with liquid N_2_ and stored at -80 °C for further analysis.

No biopsies were taken before or during the perfusion of the five human kidneys that were cultured for a 4-day period. From all eight human kidneys cultured within our platform wedge biopsies were taken at the end of culture and fixed in 4% paraformaldehyde or snap frozen in liquid nitrogen and stored at -80°C. For two of the five human kidneys that were cultured for a 4-day period we received the contralateral kidney from which control wedge biopsies were taken (Control_1 and Day4_1, Control_2 and Day4_2 in Figure 6K and Figure S5A).

### Histology

Biopsies were fixed overnight in 4% paraformaldehyde, stored in 70% ethanol and embedded in paraffin for subsequent sectioning. Periodic Acid-Schiff (PAS) staining was performed on 4-μm-thick paraffin embedded cortical tissue sections by the Pathology department using an automatic slide stainer. Tissue slices that are compared were processed simultaneously. Bright field images were digitized using a 3D Histech Pannoramic MIDI Scanner (Sysmex) and viewed with CaseViewer software. PAS stained sections of the five four-day cultured kidneys were assessed by a renal pathologist and scored on globally sclerosed glomeruli (GSG), Interstitial Fibrosis and Tubular Atrophy (IFTA), tubular cast formation, and edema.

### Tissue preparation and matrix deposition

Cryo preserved tissue biopsies were embedded in 10% gelatin and cryosectioned into 10-µm-thick sections using a Cryostar NX70 cryostat (Thermo Fisher Scientific) at –20 °C. The sections were thaw-mounted onto indium-tin-oxide (ITO)-coated glass slides (VisionTek Systems) and stored at –80 °C until further use. Slides were placed in a vacuum freeze-dryer for 15 minutes prior to matrix application. After drying, *N*-(1-naphthyl) ethylenediamine dihydrochloride (NEDC) (Sigma-Aldrich, UK) MALDI-matrix solution of 7 mg/mL in methanol/acetonitrile/deionized water (70/25/5% v/v/v) was applied using a SunCollect sprayer (SunChrom GmbH). A total of 21 matrix layers was applied at the following flow rates: layers 1–3 at 5 µL/min, layers 4–6 at 10 µL/min, layers 7–9 at 15 µL/min and layers 10–21 at 20 µL/min (speed x, medium 1; speed y, medium 1; z position, 35 mm; line distance, 1 mm; N_2_ gas pressure, 35 psi).

### MALDI-MSI measurement

MALDI-TOF/TOF-MSI was performed using a RapifleX MALDI-TOF/TOF system (Bruker Daltonics). Negative-ion-mode mass spectra were acquired at a pixel size of 5×5 µm^2^ or 20×20 µm^2^ over a mass range of *m/z* 80–1000. Prior to analysis, the instrument was externally calibrated using red phosphorus. Spectra were acquired with 15 laser shots per pixel (for 5×5 µm^2^ measurements) or 200 laser shots per pixel (for 20×20 µm^2^ measurements) at a laser repetition rate of 10 kHz. Data acquisition was performed using flexControl (Version 4.0, Bruker Daltonics) and flexImaging 5.0 (Bruker Daltonics). Sections present on the same slide were measured in a randomized order. The *m/z* features present in MALDI-TOF-MSI dataset were further used for identity assignment of metabolites and lipid species. The *m/z* values were imported into the Human Metabolome Database [24] (https://hmdb.ca/) after re-calibration in mMass and annotated for metabolites and lipids species with an error ≤ ±20 ppm. The ^13^C-labeled peaks were selected by comparing the spectrum of control and ^13^C-labeling experiments, and annotated based on the presence of un-labeled metabolites and their theoretical *m/z* values. Peak intensities of the selected features were exported for all the measured pixels from SCiLS Lab 2016b (version 2016b, Bruker Daltonics), which were used for the following analysis. Single ion visualizations were also obtained from SCiLS Lab.

### Post-MALDI-MSI staining

Following the MALDI-MSI data acquisition, excess matrix was removed by washing the slides in 100% ethanol (2×5 min), 75% ethanol (1×5 min), and 50% ethanol (1×5 min), after which tissues on the slide were fixed using 4% paraformaldehyde for 10 minutes. Slides were either stained with Mayers Haematoxylin (109249, Merck) and Eosin (E6003, Sigma-Aldrich) for brightfield imaging or with immunofluorescent antibodies. For immunofluorescent staining, slides were blocked with 3% normal donkey serum, 2% BSA, and 0.01% Triton X-100 in PBS for 1 hour at room temperature. Primary anti-CDH1 antibody (1:300, BD Biosciences, 610181), anti-nephrin (5 µg/mL, R&D, AF4269), and lotus tetragonolobus lectin (LTL, 1:300, Vector laboratories, B1325) were incubated overnight at 4 °C, followed by correspondent fluorescent-labelled secondary antibodies for 1 hour at room temperature. Slides were embedded in Prolong gold antifade mountant with DAPI (Thermo Fisher Scientific, P36931). The stained tissues were scanned using a digital slide scanner (3D Histech Pannoramic MIDI Scanner, Sysmex). Digital scanned images were aligned with the MALDI-MSI data.

### MSI data processing and analysis

For lipid analysis, features with *m/z* ≥ 400, predominately glycerophospholipids, that did not co-localize with MALDI matrix signals were selected (signal-to-noise-ratio ≥ 3). The per-pixel total ion count (TIC)-normalized intensity values for each *m/z* feature from all MSI measurements were directly exported as comma-separated values (.csv format). Upon loading in R (v. 4.0), these values were transformed into a count matrix for UMAP analysis by multiplying the intensities by 100 and taking the integer. This count data matrix was normalized and scaled using SCTransform to generate a 2-dimensional UMAP projection using Seurat (54). The spatial reconstructions of the segmentation clusters were compared to the aligned immunofluorescence stainings, and cell types were identified based on both immunomarker staining and tissue morphology. Same cell types were annotated within one cluster. The differential abundance of lipids between clusters were analyzed using the FindAllMarkers function in Seurat. To compare pixels from different datasets, these matrices were imported into the Seurat package and a data integration step was performed after batch correction using the method provided by Seurat. These integrated datasets were used to generate a 2-dimensional UMAP projection using Seurat and 3-dimensional UMAP projection using the Seurat and plotly packages. The embedding information of the 3-dimensional UMAP was translated to RGB color coding by varying red, green and blue intensities on the 3 independent axes. Together with pixel coordinate information exported from SCiLS Lab, a MxNx3 matrix was generated and used to generate molecular histology images in Matlab (v. R2019a.; Mathworks).

For dynamic metabolomics, the lipid *m/z* features from the control kidneys were used as query. Then the MALDI-MSI data from ^13^C-labeling experiments of Day_0, Day_4 and Day_8 human kidneys were used as a reference to transfer metabolite production into the query using FindTransferAnchors and TransferData function from Seurat package. Both the query and reference were normalized and scaled using SCTransform. Ultimately, all the imputed metabolite productions were combined into control kidney dataset, which contained the ^13^C-labeling information from different nutrients. The ^13^C-labeled metabolite abundance was corrected to its isotope tracer purity. Natural isotope abundance correction was performed for metabolites using R package IsoCorrectoR (55). After combining all the imputed data into one dataset and correcting the natural isotope abundance, the fraction enrichment of isotopologues was calculated based on the ratio of each ^13^C-labeled metabolite (isotopologue) to the sum of this metabolite abundance in each pixel. The calculated fraction enrichment of isotopologues was used to generate pseudo-images together with pixel coordinate information exported from SCiLS Lab. The average fraction enrichment values of identified clusters were used for generating graphs and statistical analysis. Hotspot removal (high quantile 99%) were applied to all the pseudo-images generated from calculated values.

The fraction enrichment of isotopologues derived from ^13^C_5_-glutamine were further used for relative flux rate calculation according to the previous published Q-Flux equations (44). By comparing the glutamate (m + 5) enrichment and glutamine (m + 5) enrichment, the rate of GLS relative to OGDH (V_GLS_/V_OGDH_) was determined by Equation 1. Next, comparing the two fractional contributions of aKG derived from glutamine and IDH flux yields the rate of GLS relative to IDH (V_GLS_/V_IDH_) (Equation2). The fraction of succinate derived from aKG/glu defines V_OGDH_/V_SDH(F)_ (Equation3). The fraction of malate derived directly from SDH forward flux was determined by dividing malate (m+4) enrichment by succinate (m + 4) enrichment (Equation4).

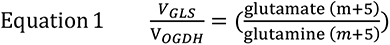

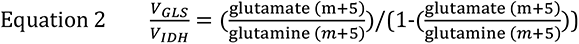

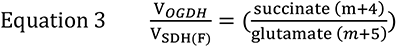

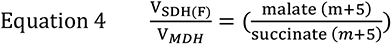

### Untargeted lipidomics

#### Sample preparation for kidney lipid extraction

Frozen kidney tissues were weighted and homogenized in cold LC-MS-grade water (final concentration: 10 mg/mL) with a Bullet Blender™ 24 (Next Advance Inc., NY, USA). Lipid extraction took place by adding 600 µL methyl-tert-butyl ether and 150 µL of methanol to 25 µL of tissue homogenate. Samples were vortexed and were kept at room temperature for 30 min. After centrifugation, the supernatant was transferred to a new microtube. The extraction was repeated with 300 µL methyl-tert-butyl ether and 100 µL methanol after which the supernatants were combined and 300 µL of water was added before centrifugating once again. The upper (non-polar) phase was transferred to glass vials and was dried under a gentle stream of N_2_. Samples were reconstituted in 100 µL 2-propanol, sonicated for 5 min and 100 µL water was added and sonicated once again for 5 min. Finally, samples were transferred to micro-vial inserts and placed in the autosampler for LC-MS/MS analysis. Quality controls (QC) were made by combining 15 µL from each sample.

#### Untargeted lipidomic analysis

Kidney lipid extracts were analysed using a LC-MS/MS based untargeted lipid profiling method (56). The LC system was a Shimadzu Nexera X2 (consisting of two LC30AD pumps, a SIL30AC autosampler, a CTO20AC column oven and a CBM20A controller) (Shimadzu, ‘s Hertogenbosch, The Netherlands). The mobile phases consisted of water:acetonitrile 80:20 (eluent A) and water:2-propanol:acetonitrile 1:90:9 (eluent B), both eluents containing 5 mM ammonium formate and 0.05% formic acid. The following gradient was applied using a flow rate of 300 µL/min: 0 min 40% B, 10 min 100% B, 12 min 100% B. A Phenomenex Kinetex C18, 2.7 µm particles, 50 × 2.1 mm (Phenomenex, Utrecht, The Netherlands) column with a Phenomenex SecurityGuard Ultra C8, 2.7 µm, 5 × 2.1 mm cartridge (Phenomenex, Utrecht, The Netherlands) as guard column were used. The column was kept at 50 °C at all times and the injection volume was 10 µL. The MS system was a Sciex TripleTOF 6600 (AB Sciex Netherlands B.V., Nieuwerkerk aan den IJssel, The Netherlands) operated in negative ESI mode (ESI-) using the following parameters: ion source gas 1 45 psi, ion source gas 2 50 psi, curtain gas 35 psi, temperature 350 °C, acquisition range *m/*z 100-1800, ion spray Voltage -4500 V (ESI-), declustering potential -80 V (ESI-). Information dependent acquisition (IDA) was used to identify lipids, with the following conditions for MS analysis: collision energy -10 V and acquisition time 250 ms; for MS/MS analysis: collision energy -45 V, collision energy spread 25, ion release delay 30, ion release width 14 and acquisition time 40 ms. The IDA switching criteria were set for ions greater than *m/z* 300 that exceed 200 cps, excluding former target for 2 s, and excluding isotopes within 1.5 Da resulting in maximum 20 candidate ions. MS-DIAL (v5.1) was used to align the data and identify the different lipids (57–59)

### Kidney auto-transplantation

Methods are shown in the supplementary methods section.

### Statistical analyses

Perfusion parameters are presented as mean ± SEM, unless indicated otherwise. MALDI-MSI derived data are presented as mean ± SD, unless indicated otherwise. Data normality and equal variances were tested using the Shapiro–Wilk test. All statistical tests were performed using GraphPad Prism 9. P-value < 0.05 were considered statistically significant. Exact sample size (*n*) for each experimental group and units of measurements were provided in the text and figure captions. All the figures were created using Adobe Illustrator (Adobe Systems).

## Acknowledgements

We would like to thank Sebastien Bouyé and Audrey Quenon (Centre Hospitalier Universitaire de Lille, Lille, France) for their expertise and assistance during the auto transplantation experiment. We would like to thank Anne Krarup Keller (Aarhus University, Aarhus, Denmark) for providing guidance in surgical techniques for the porcine auto transplantation experiment. We thank Manon Zuurmond (LUMC, Leiden, the Netherlands) for making the illustrations.

## Author contributions

Conceptualization: MdH, FW, GW, ME, TR. Data curation: MdH, FW, GW, ESL, SK, MG, BH, GW. Formal analysis: MdH, FW, ESL, SK, MG, JK, BH, GW. Funding acquisition: TR; Investigation: MdH, FW, AdG, ESL, SK, MM, TH, DdV, BH, GW, ME. Methodology: MdH, FW, SK, MG, MM, TH, DdV,BH, GW, ME, TR. Project administration: MdH, FW, GW. Resources: SK, MG, MM, TH, JKC, FP, BH. Software: SK, MG, BH, GW. Supervision: GW, ME, TR. Visualization: MdH, FW, GW. Writing – original draft: MdH, GW, TR. Writing – review and editing: MdH, FW, MJ, SK, MG, MM, DdV, JK, IA, CvK, BH, GW, ME, TR.

