## Supplemental information for "Advancing multi-day ex vivo kidney perfusion using spatially resolved metabolomics"

**LIST OF SUPPLEMENTARY MATERIALS**

**Supplementary Materials and Methods
Table S1 |** Perfusate composition **Table S2 |** Donor data of discarded human kidneys.
**Figure S1 |** Temperature-dependence of renal oxygen uptake.
**Figure S2 |** Culture platform development using porcine kidneys. Effect of continuous hemofiltration.  **Figure S3 |** Lipid heterogeneity of human kidney.
**Figure S4 |** Oxidized lipid species in perfusate during 8-day perfusion
**Figure S5 |** Histology after 4-day ex vivo perfusion of human kidneys.
**Figure S6 |** Renal function during 8-day perfusion. **Figure S7 |** Porcine kidney auto-transplantation following 4-days ex vivo culture.

**References** 60-61 (numbers for references only cited in supplementary materials)

**Supplementary Materials and Methods**

***Porcine kidney procurement***.

Kidneys from female landrace pigs (±6 months old, ±80 kg) were obtained from a local abattoir. No animal ethical committee approval was required. Pigs were sedated with an electric shock followed by exsanguination. After a warm ischemic period of 20-30 minutes kidneys were obtained and flushed with 100-200 ml of cold Ringers acetate (B. Braun) supplemented with 12.500 IE L^-1^ heparin (LEO Pharma A/S), 10 mg L^-1^ butylscopolaminebromide (Boehringer Ingelheim) and 1 mg L^-1^ nitroglycerine (Hameln). Next, kidneys were flushed with 100 ml HTK (Histidine-Tryptophan-Ketoglutarate, Custodiol) preservation solution and transported back to the laboratory on ice resulting in a cold ischemic time of 3-6 hours.

***Temperature-dependence of renal oxygen uptake***.

Porcine kidneys (n=6) were procured as stated above. During back table preparation the renal artery, vein and ureter were cannulated. The artery was flushed with cold DMEM F12, after which the kidney was placed within the organ chamber and connected to a closed-loop perfusion system. A centrifugal pump (Masterflex L/S Digital Drive 600 rpm) perfused the renal artery through silicone tubing (LS25, Masterflex Metrohm) at a mean arterial pressure (MAP) of 75 mmHg. The kidneys were perfused with the same perfusate as described in Table S1. The oxygenator was oxygenated with a carbogen mixture of 95% O_2_ and 5% CO_2_. Temperature of the perfusion fluid was controlled by a water bath connected to the oxygenator. Arterial and venous partial oxygen pressure (pO­_2_) were continuously measured using an in-line blood gas sensor (CDI 500 system, Terumo Cardiovascular Systems). Oxygen uptake (mL O_2_ min^-1^ 100 gr^-1^) was calculated as $\Delta$pO_2_ (mmHg) x Solubility of O_2_ (0.0031 mL O_2_ dL fluid^-1^ mmHg^-1^) x Renal Flow (dL min^-1^ 100 gr^-1^).

The initial temperature of the perfusion solution was set at 20˚C for a minimum of 2 hours after which the perfusate was replaced. Subsequently, baseline perfusion parameters were recorded and temperature was gradually increased. Initially to 25˚C, followed with 3˚C increments every 30 minutes until a temperature of 37 ˚C was reached.

***Effect of continuous hemofiltration***.

Porcine kidneys were procured as described above and perfused at 25°C in the platform as illustrated in Figure 1 with or without the addition of continuous hemofiltration (n=6 and n=3, respectively). Continuous hemofiltration was performed at 40 mL/h. Substitution solution was added post-hemofilter by the Prismaflex system. The substitution solution contained the same components as the culture perfusate except for Human Serum Albumin (for composition see Table S1). For both conditions, additional glucose (1M solution; D-Glucose, G8270, Sigma) was supplemented when perfusate glucose levels dropped below 4 mmol L^-1^, with the goal of maintaining euglycemia. An iSTAT1 blood analyzer (Abbott) was used for blood-gas (pH, pO_2_, pCO_2_), electrolyte (Sodium, Potassium, Chloride) and metabolite (Glucose, Lactate, Urea) measurements.

**Targeted oxidized lipid analysis**

Oxylipids were analyzed from 100 µL perfusate samples according to published protocols (60). Briefly, to 100 µL perfusate was added 100 µL water, 600 µL methanol and an internal standard mix. Following acidification with formic acid, samples were cleaned up by solid phase extraction using C18 cartridges before being analyzed by LC-MS/MS in MRM mode on a Shimadzu Nexera series UHPLC system coupled to a Sciex 6500 QTrap.

***Proof-of-concept auto transplantation***

To demonstrate that it is feasible to transplant a kidney after 4-days of ex vivo perfusion, one healthy female 40-kg mini pig (1 yr old) was used for this study. Ethical approval for a proof-of-concept auto-transplantation experiment was provided by the animal ethical review board of Lille university (n°APAFIS #31986-2021052717525766). The nephrectomy, 4-day ex vivo perfusion and auto-transplantation were performed at the animal facility of the University of Lille. To allow for acclimatization, the mini-pig arrived at a specific pathogen-free housing facility 2 weeks before the first surgery. The pig was housed, premedicated, anesthetized and monitored during surgeries according to a previously established protocol (61).

In brief, a left-sided nephrectomy was performed on Day1 of the experimental protocol. A midline incision was made after which an extraperitoneal approach was taken to mobilize the renal artery, vein and ureter. Due to splenic hemorrhage during the surgery, the peritoneal cavity was opened and a total splenectomy was performed after which the peritoneum was closed again. Following kidney procurement, the graft was immediately flushed with cold perfusate. The renal artery, vein and ureter were canulated after which the kidney was connected to the sub normothermic perfusion platform. The 4-day ex vivo perfusion was performed as described for the diseased human kidneys. On Day5, the median incision was re-opened and the kidney was transplanted into the right groin. The renal vein was anastomosed end-to-side to the caval vein and the renal artery was anastomosed end-to-side on the distal aorta. Following vascular connection, the graft was reperfused and monitored for a 3 hour period. The ureter was cannulated, and urine was collected. Assessment during this early phase of in vivo reperfusion included macroscopic appearance, doppler ultrasound of the renal blood flow and histology.

**Table S1 | Perfusate composition.**

| **Culture perfusate preparation** | | | | |
| --- | --- | --- | --- | --- |
|  | **Stock[c]** | **Final[c]** | **Per ±1L** | **Comments** |
| DMEM F12, HEPES | N/A | N/A | 700 mL | Cat# 11330, Gibco. |
| Human Serum Albumin | 200 gr L^-1^ | 20 gr L^-1^ | 100 mL | Alburex 20, CSL Behring bv. |
| Insulin-Transferrin-Sodium Selenite (100x) | N/A | N/A | 10 mL | Cat# I1884, Sigma-Aldrich.  Dissolve 1 vial in 50 mL sterile water. |
| Sodium bicarbonate | 7.5% | N/A | 10 mL | Cat# 25080094, Gibco. |
| Penicillin-streptomycin | N/A | 1% | 10 mL | Cat# 15070063, Gibco. |
| Ciprofloxacin | 2 mg mL^-1^ | 6 µg mL^-1^ | 3 mL | Fresenius Kabi. |
| Fungizone | 100 mg mL^-1^ | 0.1% | 1 mL | Bristol-Myers Squibb. |
| Citric Acid | 1M | 5 mM | 5 mL | Cat# 3200-1KG, Calbiochem. Merck.  Stock solution (1M) made by dissolving Citric Acid into sterile water. |
| Acetic Acid | 2M | 2.5 mM | 1.25 mL | Cat# 1000631000, EMSURE, Merck.  Stock solution (2M) made by Acetic Acid into sterile water. |
| Sodium Hydroxide | 1M | N/A | 20-25 mL | Cat# 567530, Calbiochem, Merck.  Stock solution (1M) made by dissolving Sodium Hydroxide pellets in sterile water. Add to perfusate until pH range is reached (7.30-7.45). |
| Sterile water | N/A | N/A | 150-200 mL | Sterile water, Versylene Fresenius.  Add to perfusate until sodium range is reached (130-145 mmol L^-1^). |
| **Substitution perfusate** | | | | |
|  | **Stock[c]** | **Final[c]** | **Per ±1L** | **Comments** |
| DMEM F12, HEPES | N/A | N/A | 700 mL | Cat# 11330, Gibco. |
| Insulin-Transferrin-Sodium Selenite (100x) | N/A | N/A | 10 mL | Cat# I1884, Sigma-Aldrich.  Dissolve 1 vial in 50 mL sterile water. |
| Sodium bicarbonate | 7.5% | N/A | 10 mL | Cat# 25080094, Gibco. |
| Penicillin-streptomycin | N/A | N/A | 10 mL | Cat# 15070063, Gibco. |
| Ciprofloxacin | 2 mg mL^-1^ | 6 µg mL^-1^ | 3 mL | Fresenius Kabi. |
| Fungizone | 100 mg mL^-1^ | N/A | 1 mL | Bristol-Myers Squibb. |
| Citric Acid | 1M | 5 mM | 5 mL | Cat# 3200-1KG, Calbiochem. Merck.  Stock solution (1M) made by dissolving Citric Acid into sterile water. |
| Acetic Acid | 2M | 2.5 mM | 1.25 mL | Cat# 1000631000, EMSURE, Merck.  Stock solution (2M) made by Acetic Acid into sterile water. |
| Sodium Hydroxide | 1M | N/A | 20-25 mL | Cat# 567530, Calbiochem, Merck.  Stock solution (1M) made by dissolving Sodium Hydroxide pellets in sterile water. Add to perfusate until pH range is reached (7.30-7.45). |
| Sterile water | N/A | N/A | 150-200 mL | Sterile water, Versylene Fresenius.  Add to perfusate until sodium range is reached (130-145 mmol L^-1^). |

**Table S2 |** **Donor data of discarded human kidneys.**

|  | **8-day Organ Culture** | | | **4-day Organ Culture** | | | | |
| --- | --- | --- | --- | --- | --- | --- | --- | --- |
|  | **Day8_1** | **Day8_2** | **Day8_3** | **Day4_1** | **Day4_2** | **Day4_3** | **Day4_4** | **Day4_5** |
| **Age (Y)** | 56 | 56 | 71 | 68 | 74 | 71 | 52 | 42 |
| **Gender (M/F)** | M | M | M | M | F | M | F | M |
| **BMI** | 30 | 30 | 25 | 24 | 27 | 25 | 16 | 16 |
| **Hypertension** | N | N | Y | N | Y | Y | N | Y |
| **Diabetes Mellitus** | N | N | Y | Y | N | Y | N | Y |
| **Smoking** | Y | Y | Y | N | N | Y | Y | Y |
| **Cause of death** | Cardiac arrest | Cardiac arrest | Cardiac arrest | CVA | SAH | CVA | Euthanasia | CVA |
| **Donor type** | DBD | DBD | DCD | DCD | DBD | DBD | DCD | DBD |
| **ICU stay (days)** | 1 | 1 | 3 | 2 | 1 | 1 | 3 | 2 |
| **Peak LDH (@ICU) (U/L)** | 943 | 943 | 760 | 239 | 684 | 282 | 196 | 206 |
| **Peak serum creatinine (@ICU) (µmol/L)** | 257 | 257 | 96 | 89 | 102 | 81 | 51 | 102 |
| **Diuresis (@ICU) (ml/hr)^1^** | 35 | 35 | 250 | 75 | 100 | 100 | 33 | 120 |
| **EGFR** | UNKN | UNKN | 82 | 83 | 82 | 95 | 122 | 99 |
| **Urine sediment protein** | 0.96 g L^-1^ | 0.96 g L^-1^ | 0.48 g L^-1^ | 0.36 g L^-1^ | < 0.20 g L^-1^ | < 0.20 g L^-1^ | < 0.20 g L^-1^ | 1.6 g L^-1^ |
| **Reason of discard** | Renal dysfunction in medical history | Renal dysfunction in medical history | Inability to allocate | Medical reasons | Medical reasons | Suspected malignancy | Inability to allocate | Previously transplanted organ |
| **Transportation** | SCS | SCS | NRP; SCS. | SCS | SCS | SCS | HMP | SCS |

**Abbreviations:** BMI, Body Mass Index; CVA, Cerebral Vascular Accident; SAH, Subarachnoid Hemorrhage; DBD, Donation after Brain Death; DCD, Donation after Cardiac Death; ICU, Intensive Care Unit; LDH, Lactate dehydrogenase; EGFR, Estimated Glomerular Filtration Function; DM, Diabetes Mellitus; SCS, Static Cold Storage; NRP, Normothermic Regional Perfusion; HMP, Hypothermic Machine Perfusion.

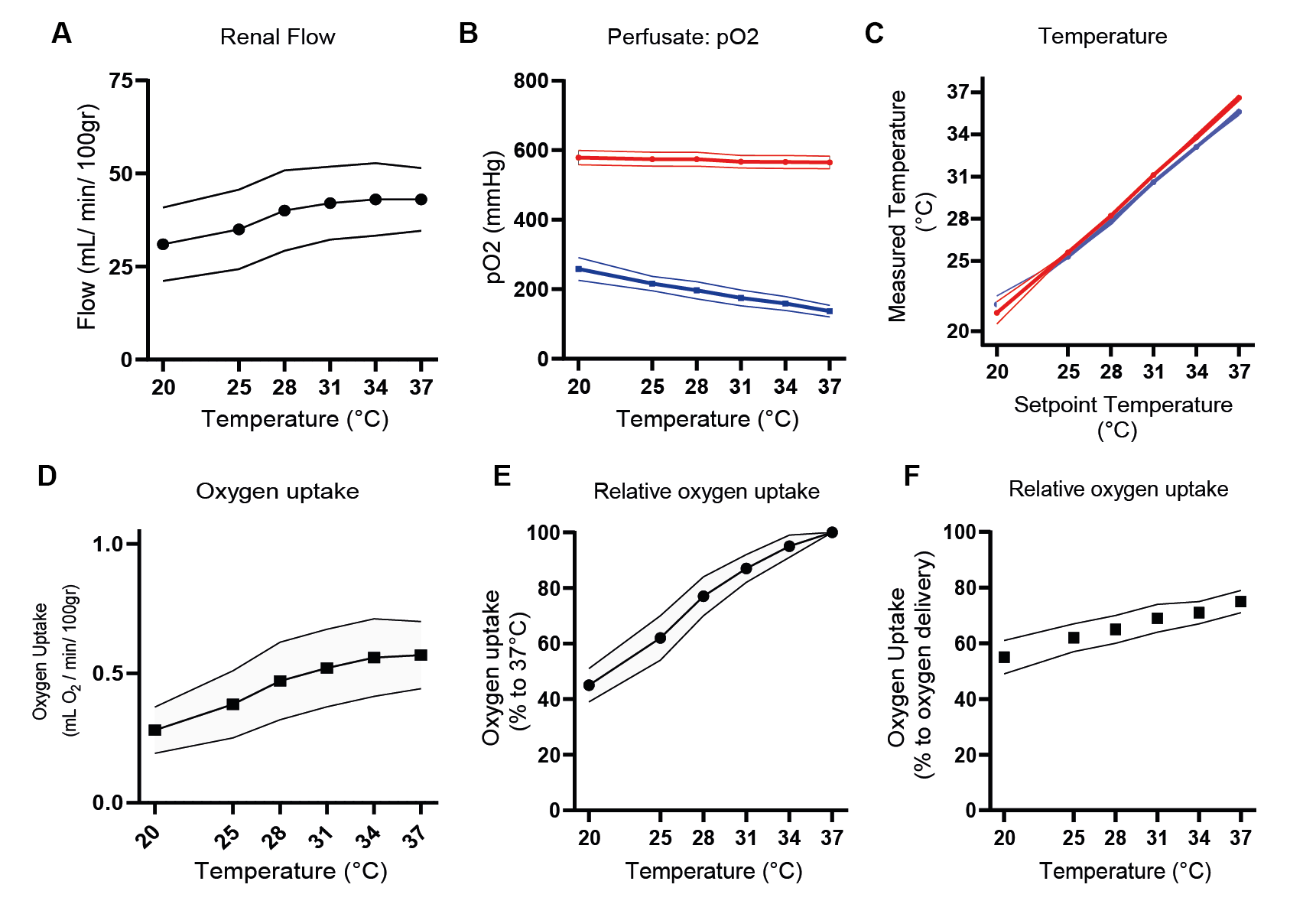

**Figure S1 |** **Temperature-dependence of renal oxygen uptake.** Temperature determines metabolic rate and therewith oxygen uptake. The effect of temperature on whole kidney oxygen uptake was determined in a closed perfusion model using porcine kidneys (n=6) that were perfused with an acellular perfusate whilst perfusion temperature was gradually increased. **A,** Average renal flow during kidney perfusion at different temperatures. **B,** Arterial and venous partial oxygen pressure during kidney perfusion at different temperatures. **C,** Measured temperature during kidney perfusion at different setpoint temperatures. **D,** Average of kidney oxygen uptake at different temperatures. **E,** Oxygen uptake relative to uptake at 37°C during kidney perfusion. **F,** Oxygen uptake relative to arterial oxygen delivery shows the percentage of available oxygen that is consumed during kidney perfusion at different temperatures. Data are presented as mean±SEM.

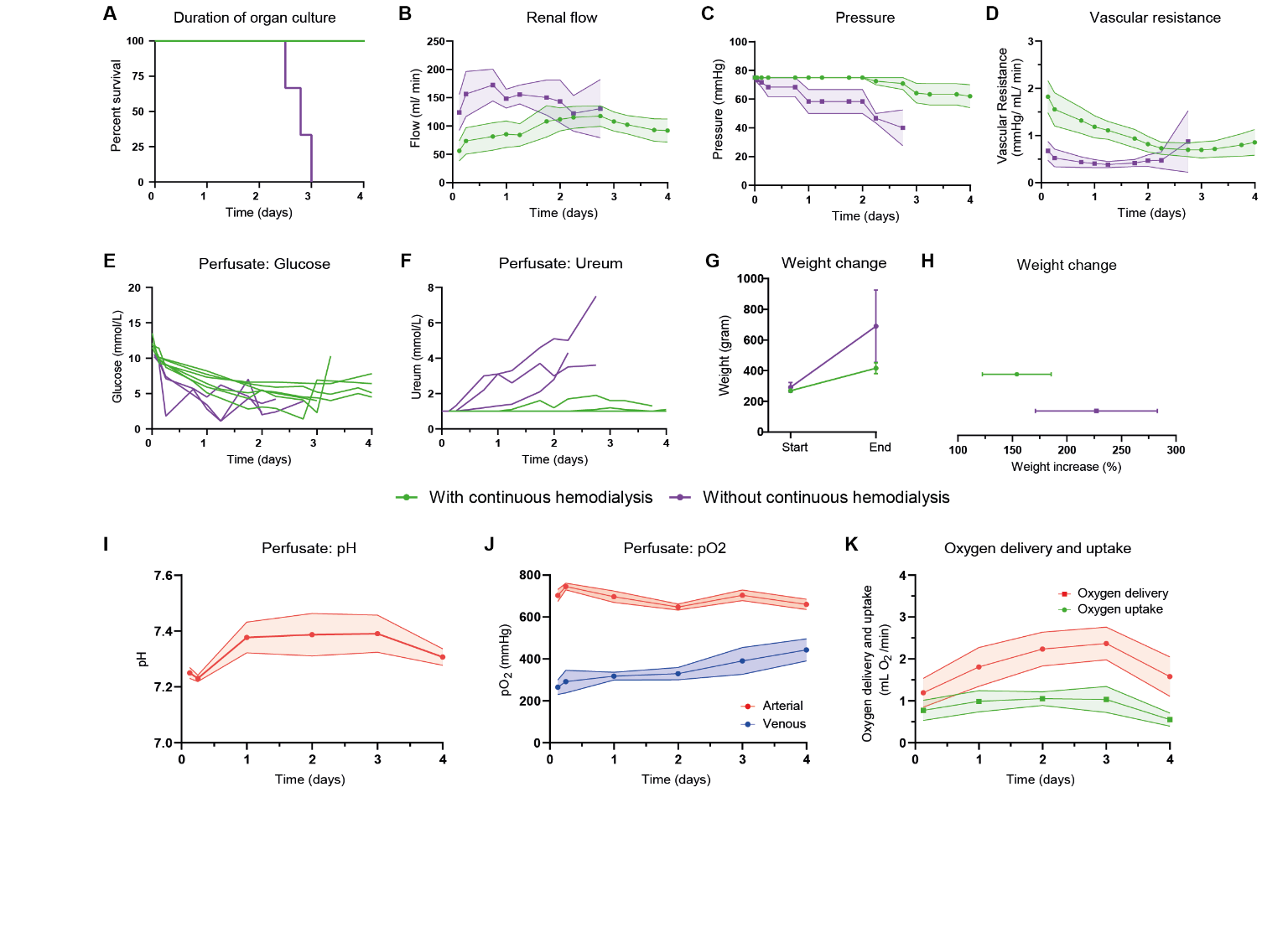

**Figure S2** **| Culture platform development using porcine kidneys. Effect of continuous hemofiltration.** Development and optimalization of sub normothermic kidney culture platform, utilizing porcine kidneys from a local abattoir. Kidneys were perfused with (n=6) and without (n=3) continuous hemodialysis. **A**, Duration of organ culture. **B-D**, Hemodynamic perfusion parameters. Renal flow (**B**), perfusion pressure (**C**) and vascular resistance (**D**) during ex vivo porcine kidney perfusion. **E**, Perfusate glucose. **F**, Perfusate ureum. **G-H**, Weight change following ex vivo perfusion with and without hemodialysis. **I-K**, Metabolic parameters of porcine kidneys cultured with continuous hemodialysis (n=6). Data are presented as mean±SEM.

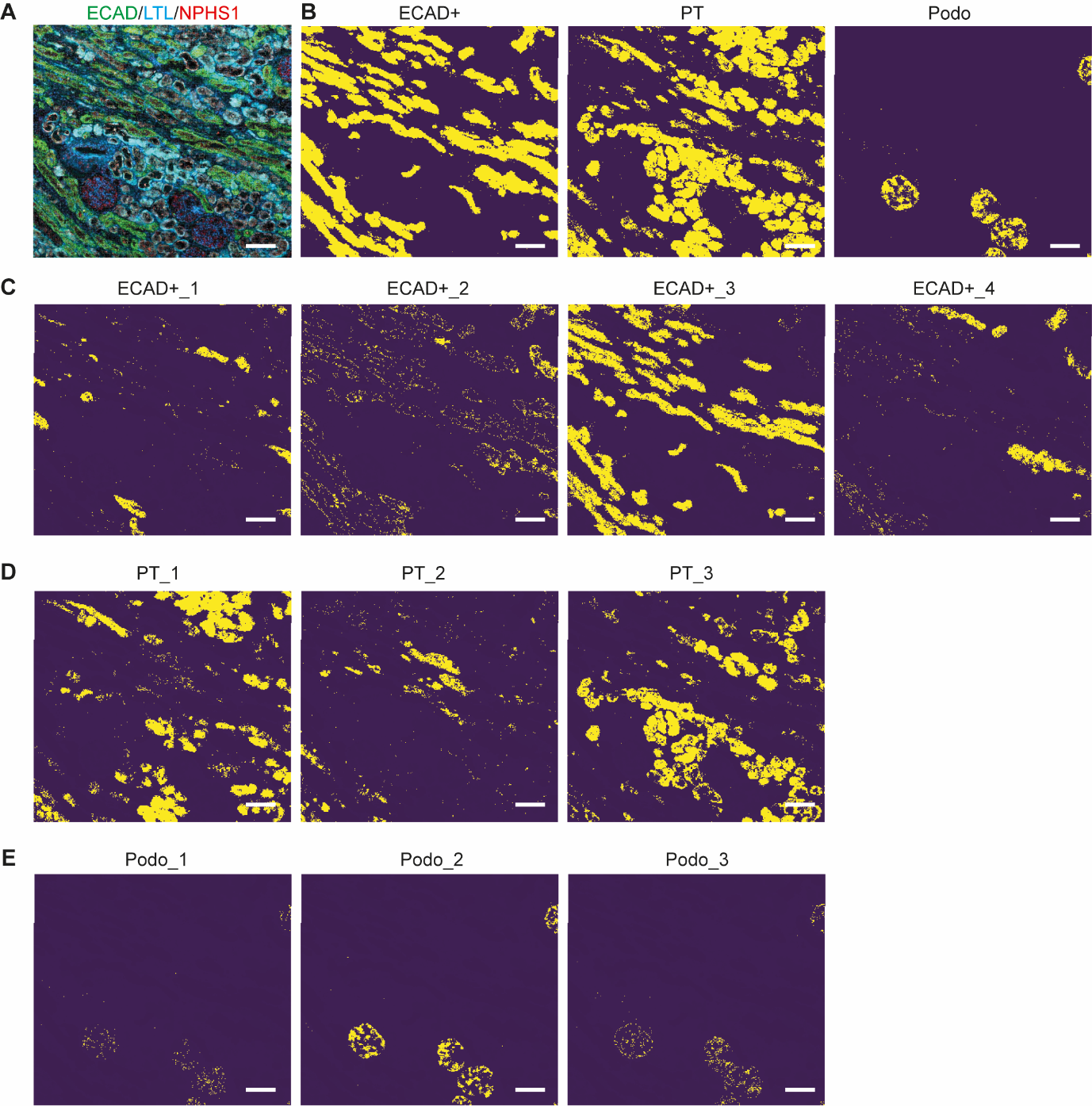

**Figure S3 | Lipid heterogeneity of human kidney.** **A,** Immunofluorescence staining (LTL, E-cadherin (ECAD) and NPHS1) on post-MALDI-MSI tissue obtained at Day8 of the 8-day organ culture period. **B,** Distribution of different epithelial cell clusters on tissue as identified in Figure 3A. **C-E,** Distribution of different phenotype of epithelial cells on tissue as identified in Figure 3B-D. Scale bars represent 200 μm.

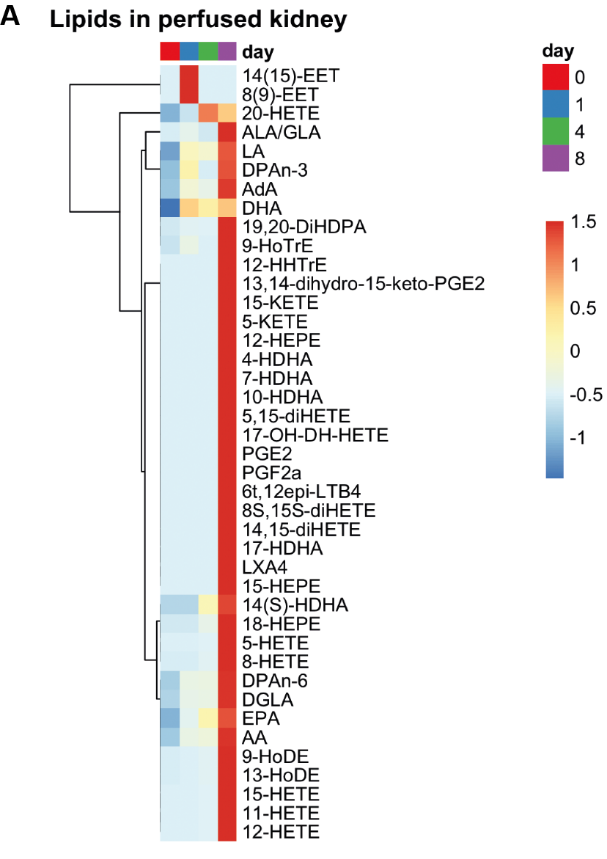

**Figure S4 | Oxidized lipid species in perfusate during 8-day perfusion**. **A**, Heatmap visualization of relative fold change in oxidized lipid species in the perfusate of Day8_2 during 8‑day perfusion, as measured by targeted lipidomics of oxylipids.

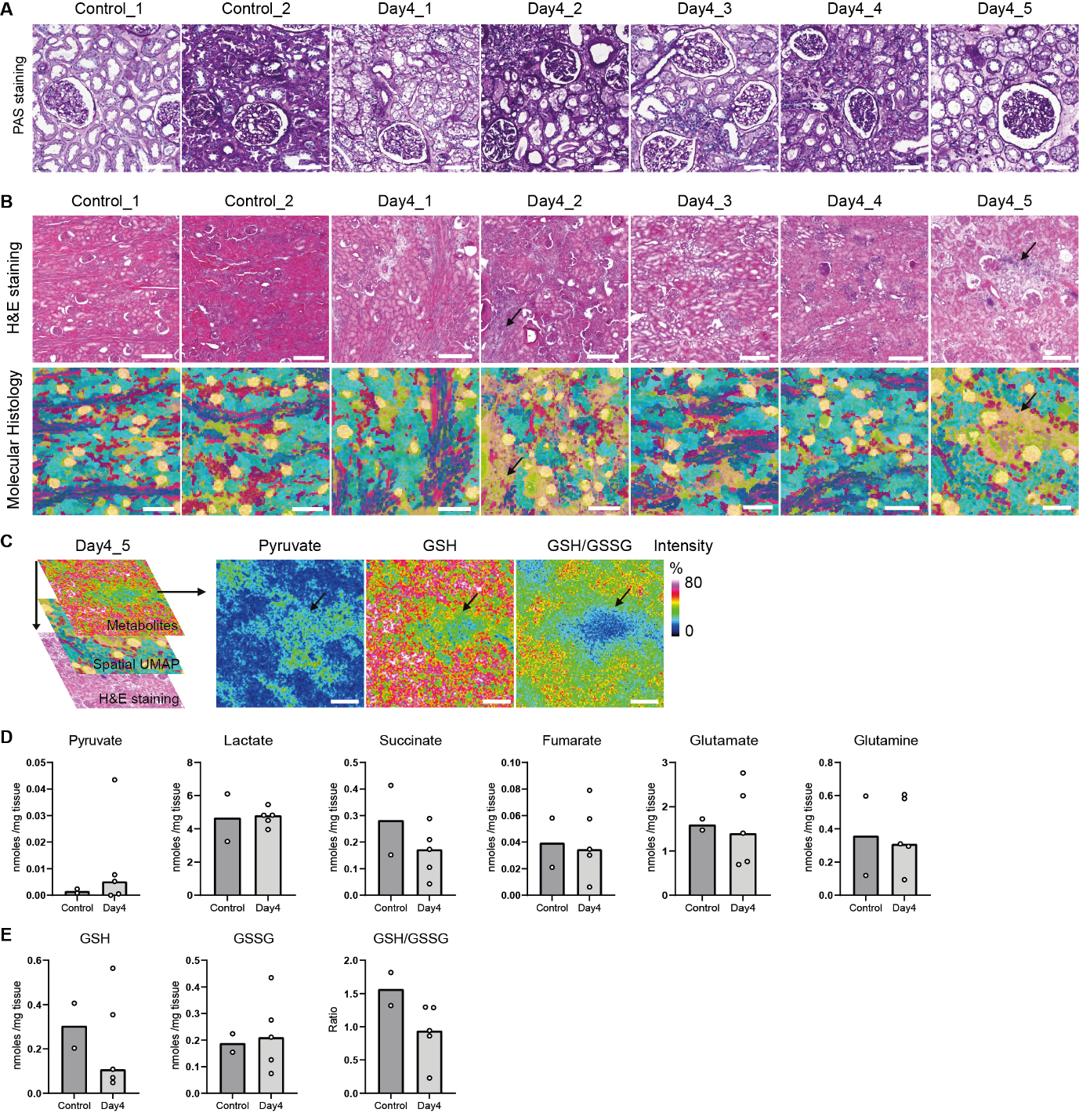

**Figure S5 | Histology after 4-day ex vivo perfusion of human kidneys.** **A**, Representative Periodic Acid-Schiff (PAS) staining for the five human kidneys that were perfused for the 4-day period. Day4_1 and Day4_2 are the contralateral kidneys of Control_1 and Control_2, respectively. Scale bars represent 100 µm.

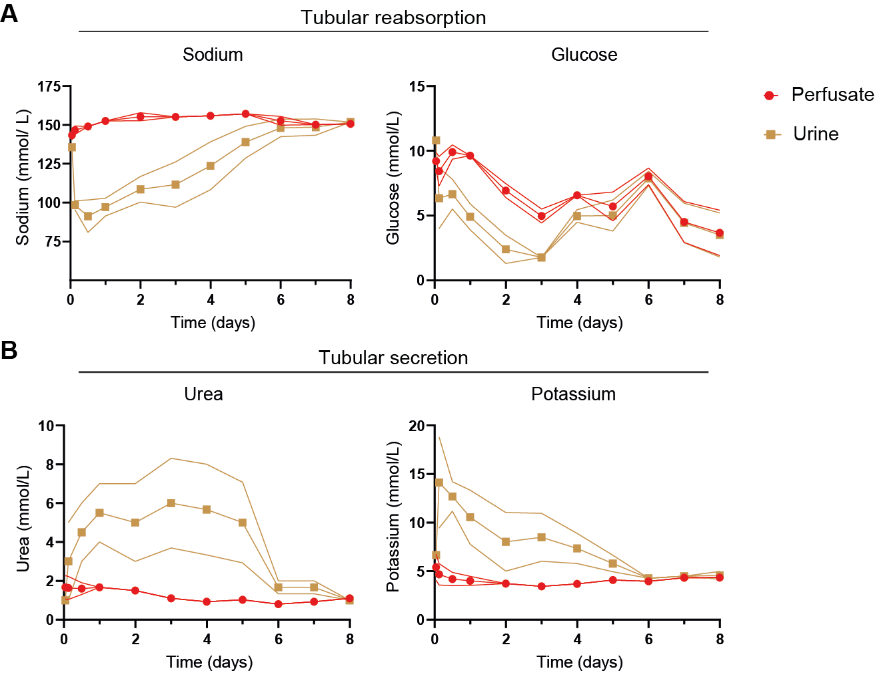

**Figure S6 |** **Renal function during 8-day perfusion.** Renal function during the 8-day sub normothermic culture of three discarded human kidneys. Concentration gradients between perfusate and urine were maintained until Day4-Day6 of perfusion. **A**, Perfusate and urine concentration of sodium and glucose demonstrates tubular reabsorption**. B**, Perfusate and urine concentration of urea and potassium demonstrates tubular secretion. Data are presented as mean±SEM.

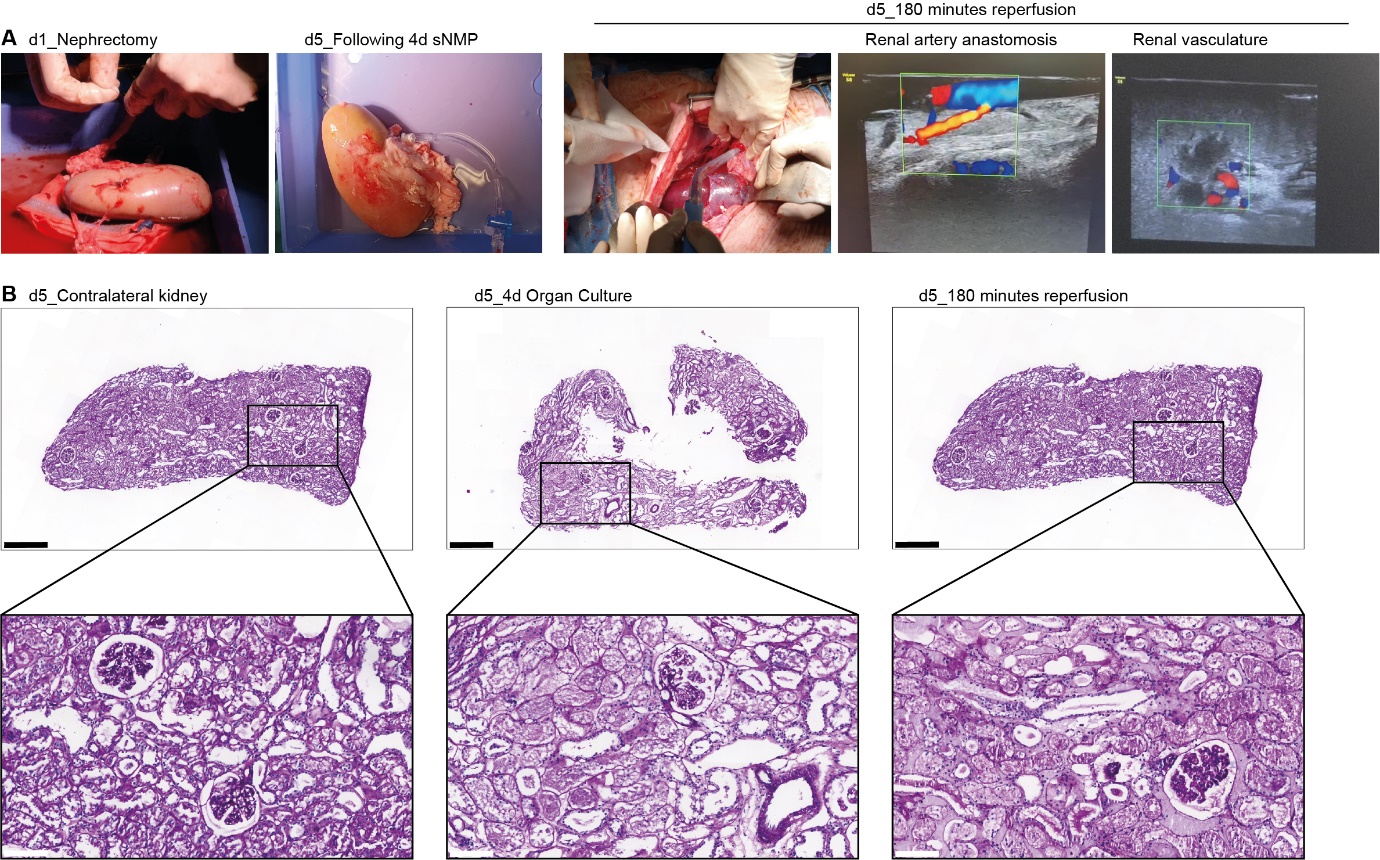

**Figure S7 |** **Porcine kidney auto-transplantation following 4-days ex vivo culture.** **A,** Macroscopic view of porcine kidney before organ culture, after 4-day organ culture, and after 180’ minutes of in vivo reperfusion. Doppler ultrasound shows perfusion through the renal artery anastomosis and within the cultured kidney 180 min after transplantation, indicative for a preserved vasculature (i.e. non-coagulant). **B**, Representative PAS staining demonstrating histology for cortical biopsies taken from the contralateral kidney on Day5, the cultured kidney on Day5 before in vivo reperfusion, and after 180 min of in vivo reperfusion. Top images: scale bars represent 400 µm. Bottom images: scale bars represent 100 µm.
